# Chaperones shape the conformational landscape of 26S-proteasome-base assembly for allosteric ATPase motor activation

**DOI:** 10.64898/2026.06.01.729410

**Authors:** Hao-Hsuan Hsieh, Andreas Martin

**Affiliations:** California Institute for Quantitative Biosciences, University of California at Berkeley, Berkeley, CA 94720, USA; Department of Molecular and Cell Biology, University of California at Berkeley, Berkeley, CA 94720, USA; Howard Hughes Medical Institute, University of California at Berkeley, Berkeley, CA 94720, USA

## Abstract

Protein homeostasis depends on the 26S proteasome, the most complex ATP-dependent protease in eukaryotic cells. The proteasome base subcomplex is responsible for mechanical substrate unfolding and translocation into an internal degradation chamber. It contains three non-ATPase subunits, Rpn1, Rpn2, and Rpn13, and a heterohexameric AAA+ motor with six distinct ATPases, Rpt1 – Rpt6. Correct base assembly requires four dedicated chaperones that initially form the Hsm3 module (Hsm3-Rpt1-Rpt2-Rpn1), the Rpn14/Nas6 module (Rpn14-Rpt6-Nas6-Rpt3-Rpn2-Rpn13), and the Nas2 module (Nas2-Rpt5-Rpt4). However, the mechanisms underlying module assembly and formation of the mature base remain unknown. Here, we *in vitro* reconstitute the base subcomplex of the *S. cerevisiae* 26S proteasome from recombinant modules. Using biochemical assays, mass photometry, single-molecule fluorescence measurements, and single-particle cryo-EM, we reveal how the chaperones direct the conformational transitions through several intermediates toward the ATP-hydrolysis-active base. The Nas2 and Rpn14/Nas6 modules associate first, and binding of the Hsm3 module creates a state in which the chaperones stabilize an open ATPase ring that lacks hydrolysis activity. Sequential chaperone release then leads to a gradual ATPase-ring closure, whereby Hsm3’s unstructured C-terminal tail mimics a substrate polypeptide in the central channel and induces a processing motor state with a spiral-staircase arrangement of Rpt subunits and a closed ATPase site at Rpt4. Inaugural ATP hydrolysis in Rpt4 is subsequently required to eject Hsm3 and transition to the Nas6-bound base that is ATPase active and competent for 26S-proteasome incorporation. Our studies thus provide exciting insights into how chaperones assure correct assembly, guide the complex through an intricate conformational landscape, and thereby prevent premature ATP-hydrolysis activation or incorporation of faulty assemblies into holoenzymes.

## Introduction

Numerous macromolecular machines that are crucial to cellular homeostasis require dedicated chaperones or assembly factors to ensure their quality and timely biogenesis. Among these molecular machines, the 26S proteasome maintains proteostasis and regulates key cellular processes by degrading misfolded, damaged, or regulatory proteins^1–3^. It is composed of two major subcomplexes, the 20S core particle (CP) and the 19S regulatory particle (RP), with the RP consisting of two subcomplexes, the lid and the base. The RP captures substrates in ubiquitin-dependent or -independent, adaptor-mediated manners, unfolds them, and translocates the unstructured polypeptides into the internal degradation chamber of the barrel-shaped CP for proteolytic cleavage into peptides. The lid with its nine subunits primarily has scaffolding functions, but also includes Rpn11 as the major deubiquitinase of the proteasome. The base subcomplex consists of three non-ATPase subunits, Rpn1, Rpn2, and Rpn13, with ubiquitin-receptor and scaffolding roles, and six distinct ATPase subunits of the AAA+ (ATPases Associated with diverse cellular Activities) family that form a heterohexameric ring in the clockwise order Rpt1, Rp2, Rpt6, Rpt3, Rpt4, and Rpt5 ^4^. Assembling the base subcomplex with the correct composition and functional arrangement of subunits that allows the subsequent formation of 26S proteasome holoenzymes thus represents a particular challenge in macromolecular complex formation.

Each Rpt subunit contains an N-terminal helix that in the fully assembled hexamer forms a coiled-coil with a neighboring Rpt, leading to a pairing of Rpt1/Rpt2, Rpt6/Rpt3 and Rpt4/Rpt5. The coiled coils of Rpt1/Rpt2 and Rpt6/Rpt3 additionally provide the binding sites for the large non-ATPase subunits Rpn1 and Rpn2, respectively. The coiled coil is followed by a N-terminal globular domain (NTD) with Oligonucleotide/Oligosaccharide-Binding (OB) fold that in the hexamer context constitutes a rigid N-ring. Underneath this N-ring, the C-terminal ATPase domains of Rpt1 - Rpt6 form the motor ring with a typical AAA+ architecture. Each ATPase domain consists of a large and a small AAA+ subdomain, with the ATP-binding site located at the interface to the clockwise-neighboring subunit. An intra-subunit Walker-B motif positions Mg^2+^ and a catalytic water molecule, while arginine finger motifs from the clockwise neighboring subunit coordinate the γ-phosphate of ATP for hydrolysis. A pore-1 loop with a highly conserved aromatic residue (Tyr in the *S. cerevisiae* 26S proteasome) projects from each ATPase domain into the central channel, sterically interacts with the substrate polypeptide, and allows substrate translocation through mechanochemical coupling in a hand-over-hand motion ^5,6^. Five ATPase domains are thereby arranged in righthanded spiral-staircase conformations, with the sixth representing the seam subunit located between the bottom and the top of the staircase.

Four dedicated chaperones, Nas2, Nas6, Rpn14, and Hsm3 in yeast (Psmd9, Psmd10, Paaf1, and Psmd5 in humans, respectively), have been identified to facilitate the assembly of the base subcomplex ^7–11^. Rpt1/Rpt2, Rpt6/Rpt3, and Rpt4/Rpt5 likely form dimers co-translationally through interactions of their N-terminal coiled-coil domain ^12^, before the Rpn subunits and chaperones bind to form three base-assembly modules: the Nas2 module (Nas2-Rpt5-Rpt4) the Rpn14/Nas6 module (Rpn14-Rpt6-Nas6-Rpt3-Rpn2-Rpn13), and the Hsm3 module (Hsm3-Rpt1-Rpt2-Rpn1) ^7,8,13–16^.

Nas2 consists of four N-terminal α-helices and a C-terminal Psd95-Dlg1-Zo1 (PDZ) domain, joined by a short flexible linker ^17,18^. The structure of the N-terminal helices has been solved alone and associated with the C-terminal small AAA+ subdomain of Rpt5, with α3 and α4 of Nas2 forming the binding interface. The PDZ-domain structure has also been solved in isolation, but not in complex with Rpt5. Members of the PDZ-domain family generally interact with the C-terminus of their binding partners, suggesting a similar interaction between Nas2’s PDZ domain and the C-terminus of Rpt5 ^19^. This model is supported by previous NMR data for a complex between fragments of Rpt5 and Nas2, and by the finding that Rpt5’s three C-terminal residues are important for Nas2 binding ^20,21^. Previous cell biology studies also suggest that Nas2 dissociation is a checkpoint during base assembly ^22^, yet the exact roles of Nas2 and its mechanisms for release remain unclear, and there are no structural data for full-length Nas2 in the context of base assembly.

The Rpn14/Nas6 module is characterized by Rpn14 interacting with Rpt6 and Nas6 binding to the C-terminus of Rpt3. The Nas6-Rpt3 interaction has been well characterized, with structural data ranging from the Nas6-bound small AAA+ subdomain fragment of Rpt3 to Nas6-bound RP ^23–25^. The consensus is that Nas6 forms a structure of seven ankyrin repeats that cradle Rpt3’s C-terminal small AAA+ domain to regulate RP-CP association ^26–28^. On the other hand, the Rpn14-Rpt6 interactions remain much more elusive, with only the structure of Rpn14 in isolation available ^29,30^. Rpn14 consists of an N-terminal domain with six β-strands and one helix, followed by a C-terminal WD40 domain with a seven-bladed β-propeller. Based on structural analyses and NMR data for a complex with an Rpt6 fragment, the current model suggests that Rpn14 interacts with Rpt6’s small AAA+ subdomain ^30,31^. However, how Rpn14 binds full length Rpt6, how it forms the Rpn14/Nas6 module, and what regulatory roles it plays during base assembly remain unknown.

Hsm3 consists of eleven HEAT repeats that form a horseshoe-shaped structure, followed by a C-terminal intrinsically disordered region (IDR) with ∼ 16 residues. Structural and mutational data indicated that it directly interacts with the C-terminal small AAA+ subdomain of Rpt1 and may also contact Rpt2 ^32,33^. Deletion of Hsm3 in yeast is the most defective of all single chaperone deletion mutants and causes base-assembly intermediates to build up in the cell ^11,22^. Recent mass spectroscopy studies showed that Hsm3 crosslinks to all Rpts, with Rpt1 and Rpt2 forming crosslinks to the structured part of Hsm3, while Rpt6, Rpt3, Rpt4, and Rpt5 crosslink to its C-terminal tail ^34^. Structural data on the Hsm3-bound RP from the same study also suggested a potential insertion of the C-terminal tail into the central channel of the Rpt ring. However, despite Hsm3’s importance, it is unclear how it chaperones Rpt1 and Rpt2, and regulates base assembly.

For the yeast proteasome, it is proposed that the Nas2 module first associates with the Rpn14/Nas6 module, before the Hsm3 module joins the complex ^4,22^. For the mammalian proteasome, however, the current model is that the Hsm3 module first associates with the Rpn14/Nas6 module instead ^8^. Despite the conservation of the base subcomplex and its chaperones from yeast to human, these conflicting models for the order of assembly steps highlight the major gaps in our understanding of base formation and proteasome biogenesis in general. While Nas6 stays associated with RP until being evicted from nascent 26S proteasomes through conformation-specific allosteric interactions ^27^, Nas2, Rpn14, and Hsm3 likely dissociate earlier during the base-assembly process and prior to formation of the RP ^11^. Based on current structural models and predictions, the modes of Nas2 and Hsm3 binding may conflict with the formation of the mature ATPase ring, suggesting that their dissociation is coupled to quality checkpoints during base assembly. However, all previous structural data were acquired with individual chaperones and RP or fragments of Rpt subunits, outside the context of base assembly. The molecular mechanisms and temporal coordination that underlie module assembly, chaperone release, and the formation of a mature, ATP-hydrolysis-competent base subcomplex are therefore outstanding questions.

Proteasomal base assembly has so far been studied primarily through knockouts and pulldowns from cell extracts to analyze the accumulation of intermediates, yet these investigations were significantly hampered by the transient nature of intermediates in the cell. In this study we therefore reconstituted the assembly of the yeast proteasomal base from recombinantly expressed modules *in vitro*, which allowed detailed structural and biophysical characterizations with precise control of components and conditions. Using mass photometry ^35,36^, we identified key intermediates during base assembly and the requirements for the release of Nas2, Rpn14, and Hsm3. We developed a single-molecule fluorescence-based assay to monitor the assembly process from reconstituted modules in real time, and we structurally investigated the intermediates by cryogenic electron microscopy (cryo-EM), which revealed the modes of chaperone interactions with individual subunits and the conformational transitions during the stepwise chaperone release. These results show how the chaperones shape the conformation landscape of the intermediate to regulate the assembly process and lead to the mature, ATP-hydrolysis active base.

## Results

### Reconstitution of base assembly *in vitro*

To study the mechanisms and structural features of the proteasomal base assembly, we recombinantly expressed the previously identified yeast *S. cerevisiae* Nas2, Rpn14/Nas6, and Hsm3 modules in *E. coli*. Following their purification by affinity and size-exclusion chromatography (Extended Data Fig. 1A), we first assessed their assembly states and interactions by mass photometry (Fig. 1). Each module showed the expected mass distribution (∼ 269 kDa for the Hsm3 module, 289 kDa for the Rpn14/Nas6 module, and 125 kDa for the Nas2 module, Fig. 1A), indicating the presence of all Rpt, Rpn, and chaperone subunits. In addition, we detected smaller species due to some extent of subunit dissociation that occurred at the low concentrations (∼ 5 nM) required for the mass photometry measurements. Pairwise combination of the modules led to complex formation only between the Nas2 and Rpn14/Nas6 modules (MW ∼ 415 kDa), whereas the Hsm3 module did not interact with either of the other two modules alone (Fig. 1A). This is consistent with previous pulldown experiments from yeast cell extracts that revealed a Nas2-Rpt5-Rpt4:Rpn14-Rpt6-Nas6-Rpt3-Rpn2-Rpn13 intermediate, but no two-module intermediates involving Hsm3 ^22^. Mixing all three modules led to a fully assembled base complex with a molecular weight of ∼ 556 kDa, which is in agreement with the *in vivo* assembled base that contains only the Nas6 chaperone (Figure 1A). Again, low amounts of partially assembled species were present due to the sub-saturating module concentrations used in these experiments. Overall, these results indicate that the Rpn14/Nas6 and Nas2 modules associate first, and the Hsm3 module joins later, leading to the subsequent release of the Nas2, Rpn14 and Hsm3 chaperones to form a Nas6-bound base complex.

**Figure 1.**
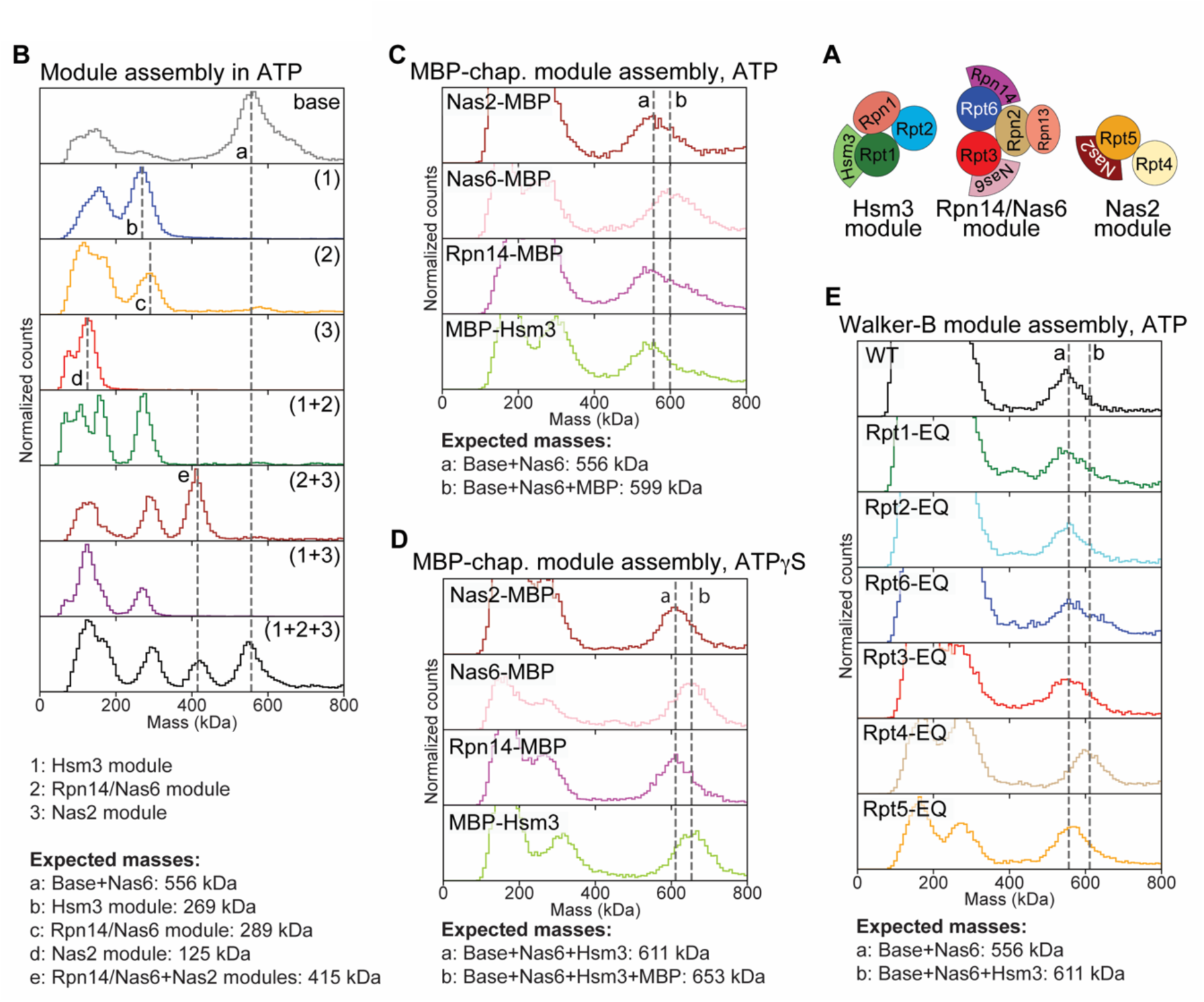
Mass-photometry measurement of base assembly modules and intermediates. (A) Schematics showing the subunit compositions of the Hsm3, Rpn14/Nas6, and Nas2 module. (B) Mass distributions of the base-assembly modules and their mixtures in ATP. The “base” sample represents the recombinant Nas6-bound base subcomplex purified from *E. coli* as previously described ^37^, with the base including the subunits Rpt1, Rpt2, Rpt3, Rpt4, Rpt5, Rpt6, Rpn1, Rpn2, and Rpn13. (C, D) Base assembly in ATP or ATPγS from modules that contained a single MBP fusion of the four chaperones. A 42.5 kDa mass shift indicates that a particular MBP-fused chaperone remained bound to the base-assembly product. While only Nas6 stays bound in the presence of ATP, both Nas6 and Hsm3 are retained in the presence of ATPγS. (E) Base assembly from modules containing individual Walker-B (EQ) mutant Rpt subunits. Only assembly with the Rpt4-EQ mutant caused a mass shift, corresponding to the retention of Hsm3. Mass-photometry counts were normalized to allow comparisons between panels, and dashed lines represent the expected mass of the complex indicated in each plot.

We confirmed the functionality of this *in vitro* assembled base by ATP-hydrolysis assays, where the complex reconstituted from all three modules exhibited the same activity as the *in vivo* assembled base with Nas6 (k_ATPase_ = 1.06 +/- 0.01 s^-1^; Extended Data Fig. 1B). In contrast, neither of the individual modules nor the intermediate consisting of the Nas2 and Rpn14/Nas6 modules exhibited considerable activity (Extended Data Fig. 1B), suggesting that ATP hydrolysis requires the presence of all six Rpt subunits and potentially the release of the Nas2, Rpn14, and Hsm3 chaperones.

Given the ATPase nature of the base, we hypothesized that the assembly and/or the dissociation of chaperones may be regulated by ATP hydrolysis. We therefore compared the products of module assembly in the presence of ATP or the slowly-hydrolyzing analogue, ATPγS. Importantly, these experiments were performed with modules in which the individual chaperones were fused to the maltose binding protein (MBP) in order to conclude from molecular-weight shifts which chaperone was potentially retained. For assembly in ATP, only the Nas6-MBP fusion showed a ∼ 42.5 kDa shift in the mass distribution, consistent with Nas6 being the only bound chaperone in the fully assembled base (Fig. 1B). Interestingly, assembly in ATPγS resulted in a complex with a molecular weight of ∼ 611 kDa, and MBP tagging led to a corresponding mass shift for both Nas6 and Hsm3 (Fig. 1C). We therefore concluded that ATP hydrolysis is required for the dissociation of Hsm3, while Nas2 and Rpn14 dissociate spontaneously upon the joining of all three modules. Since the base ATPase ring consists of six distinct Rpts that differentially contribute to its activities ^37,38^, it is possible that ATP hydrolysis in these subunits also differentially affect chaperone release. We therefore generated modules in which individual Rpts were inactivated in their ATP hydrolysis by a Glu to Gln (EQ) mutation in the Walker-B motif. Assembly of these modules in ATP revealed that Hsm3 release depends on hydrolysis by Rpt4, but is independent of hydrolysis in other Rpts (Fig. 1D). Importantly, the Rpt4-EQ mutant does not seem to cause other unintended changes, as module assembly in the presence of ATPγS looked identical for all EQ mutants and wild-type Rpts (Extended Data Fig. 1C).

### Single-molecule studies of chaperone release

To provide further insight into the order and kinetics of chaperone release during module assembly, we used single-molecule total internal reflection fluorescence (TIRF) microscopy. The Nas2 module containing LD555-labeled Nas2 and N-terminally biotinylated Rpt4 was pre-assembled with the Rpn14/Nas6 module and immobilized through neutravidin on biotin-coated PEGylated slides. We then added an Hsm3 module with LD655-labeled Hsm3 chaperone and monitored module arrival and chaperone release by following the signals for LD555 and LD655 fluorescence upon simultaneous excitation (Fig. 2A). These measurements showed that the appearance of the LD655 signal, reporting on incorporation of the Hsm3 module, is quickly followed by disappearance of the LD555 signal, which indicates the triggered release of Nas2 with a dwell time of 1_Nas2_ ∼ 0.3 +/- 0.1s. During the brief presence of both labeled chaperones, we observed weak FRET (Fig. 2B), indicating the proximity of Nas2 and Hsm3 in the base assembly intermediate. Hsm3 is then expelled, indicated by the loss of the LD655 signal, with 1_Hsm3_ ∼ 6.6 +/- 2.1 s after Hsm3 module association (Extended Data Fig. 2). Corresponding measurements for the release of Rpn14 remained unsuccessful, primarily due to sparsity of events, caused by the low affinity of modules combined with the low concentrations required for single-molecule TIRF measurements. However, based on our mass photometry experiments we can conclude that Rpn14 leaves after Nas2 and before Hsm3.

**Figure 2.**
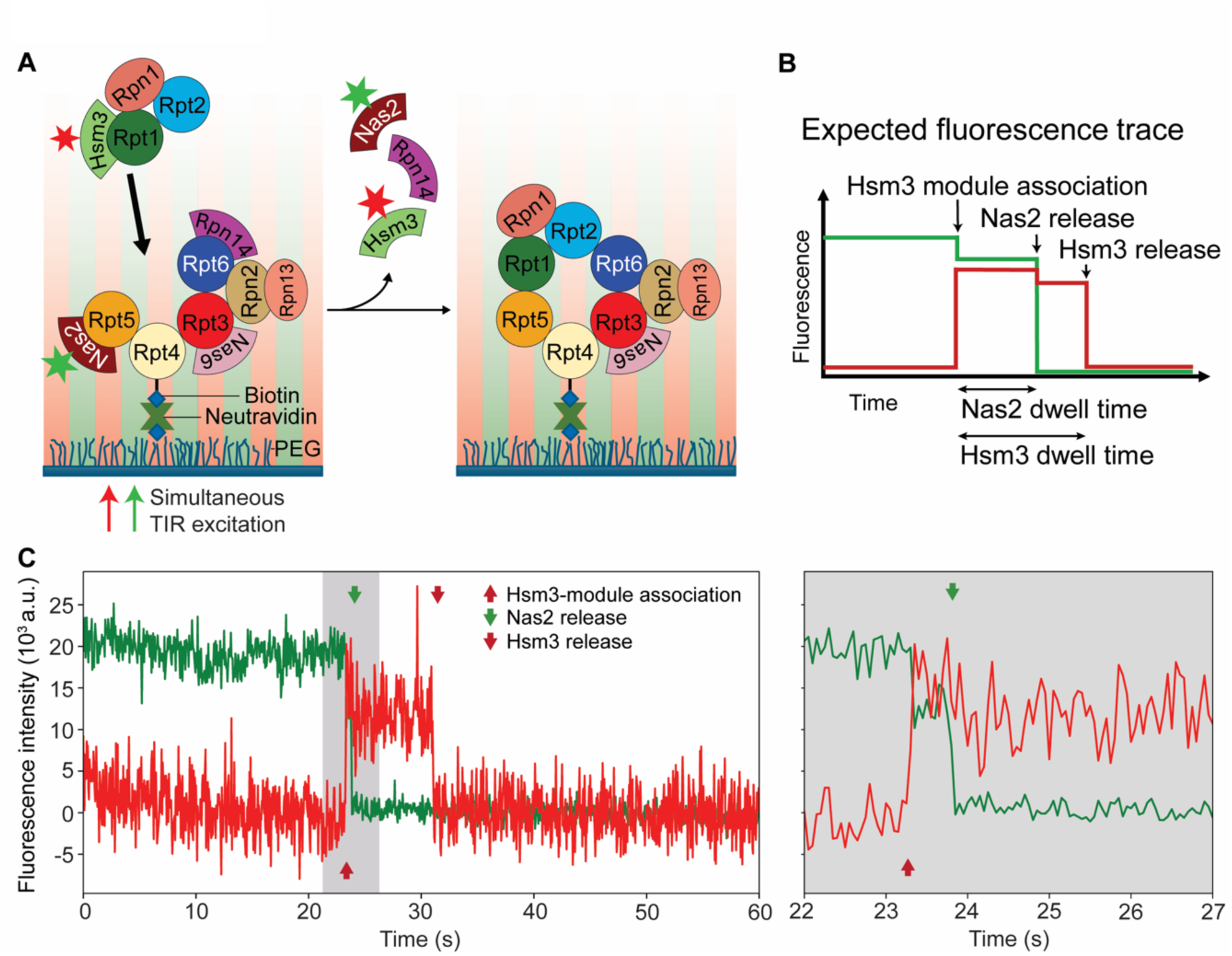
Single-molecule fluorescence measurements of Nas2 and Hsm3 release following Hsm3-module association. (A) Schematic for the single-molecule TIRF experiments, following the binding of the Hsm3 module with LD655 (red star) labeled Hsm3 to the preassembled and immobilized Rpn14/Nas6 and Nas2 modules containing LD555 (green star) labeled Nas2. Presence of Hsm3 and Nas2 were monitored through simultaneous total internal reflection (TIR) excitation and detection of their fluorescent labels. (B) Expected fluorescence time trace with intensity changes corresponding to important molecular events. The increase in red fluorescence upon association of the Hsm3 module is accompanied by a reciprocal small drop in green fluorescence due to weak FRET between Nas2 and Hsm3. Loss of this FRET after Nas2 release causes a minor drop in red fluorescence. (C) Example for an observed fluorescence time trace, shown complete (left) and zoomed-in to visualize the Nas2 release shortly after Hasm3-module arrival. The time points for Hsm3-module association, Nas2 release, and Hsm3 release are indicated by red and green arrows.

### Structures of base assembly intermediates

Our discovery that Hsm3 release depends on ATP hydrolysis in Rpt4 provided the opportunity to stabilize Hsm3-bound base-assembly intermediates for cryo-EM structure determination. We therefore prepared grids with *in vitro* reconstituted base containing the Rpt4-EQ mutant and collected a large data set whose particles could be classified into three major compositional states: base-Nas2-Rpn14-Hsm3-Nas6, base-Rpn14-Hsm3-Nas6, and base-Hsm3-Nas6 (Fig. 2A-C). Intriguingly, a significant number of particles showed density for Nas2 (∼15%) and/or Rpn14 (∼65%). This may seem unexpected based on our *in vitro* assembly data from mass photometry, but can be explained by the about three orders of magnitude difference in concentrations: ∼ 5 nM for mass photometry and ∼ 10 µM for cryo-EM. Correspondingly, a dilution of the cryo-EM sample showed the same mass distribution in mass-photometry measurements as the mass-photometry samples described above (Extended Data Fig. 1D). Thus, the dissociation of Nas2 and Rpn14 may be a reversible process before ATP hydrolysis in Rpt4, and their occupancies in the assembly intermediate are determined by their concentrations.

We refined the three conformations to nominal resolutions of 4.9 Å, 3.9 Å, and 3.8 Å for the base-Nas2-Rpn14-Hsm3-Nas6, base-Rpn14-Hsm3-Nas6, and base-Hsm3-Nas6 complexes, respectively (Extended Data Fig. 3,4; Supplemental Table 1), allowing us to build molecular models (Fig. 3 and Extended Data Fig. 5; Supplemental Video 1). All three states are highly dynamic, with Rpn1, the ATPase portions of Rpt6, Rpt3, Rpt4, and Rpt5, and most of the Rpt1-Rpt2-Hsm3 assembly moving independently relative to each other, the N-ring, and Rpn2. Despite these dynamics, Nas6 was observed in all three complexes in an identical conformation, interacting with the small AAA+ subdomain of Rpt3 and resembling a previous crystal structure of Nas6 bound to a C-terminal fragment of Rpt3 (Extended Data Fig. 6A-C; Supplemental Video 2) ^24,25^. These findings indicate that Nas6 mainly regulates processes downstream of the base assembly itself. By contrast, Nas2, Rpn14, and Hsm3 interact with several subunits, determine their conformational landscapes, and thus direct an ordered assembly and activation of the ATPase ring.

**Figure 3.**
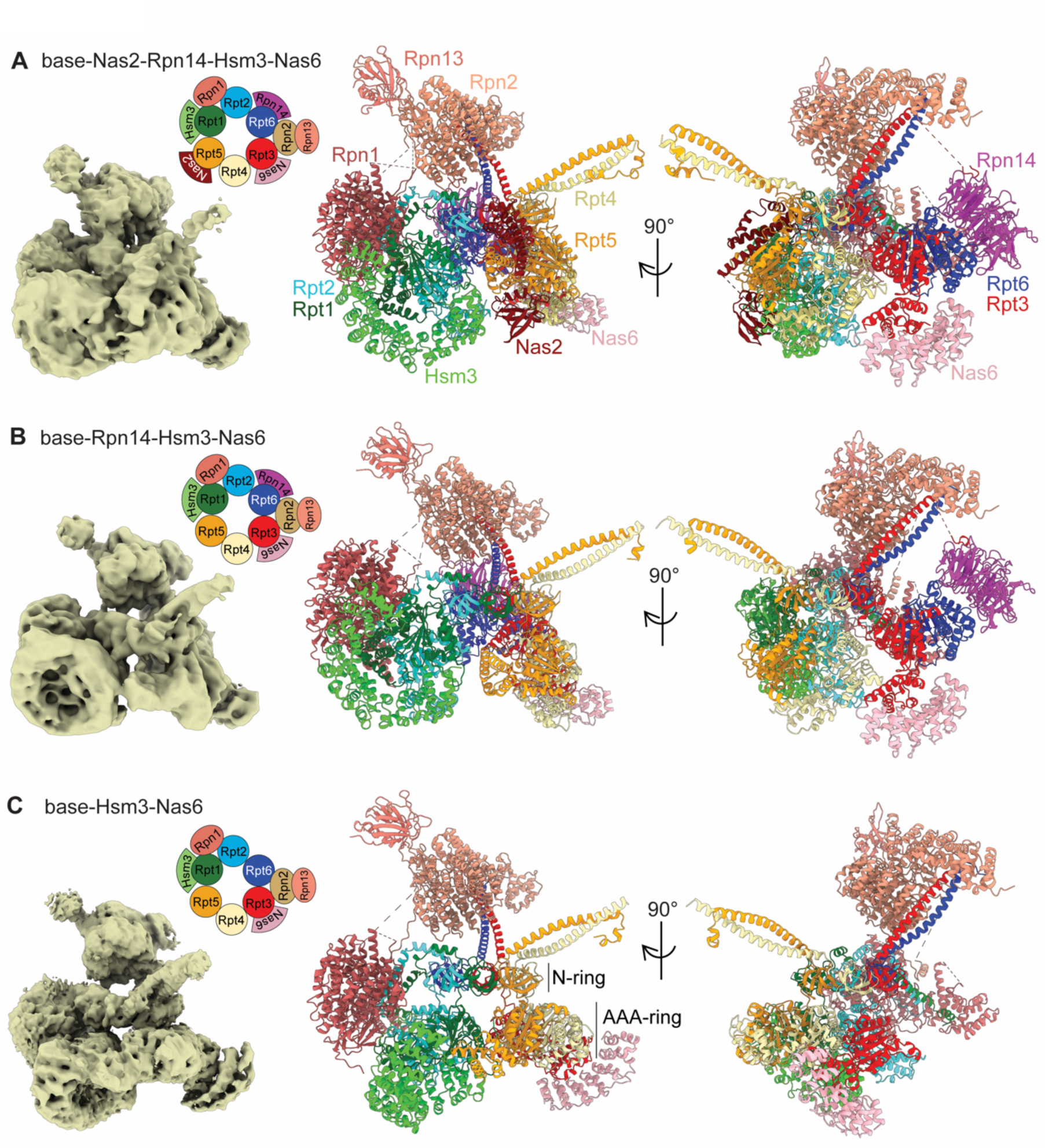
Cryo-EM structures of base assembly intermediate states. Cryo-EM density maps (left) and the corresponding molecular models (middle and right) for (A) base-Nas2-Rpn14-Hsm3-Nas6, (B) base-Rpn14-Hsm3-Nas6, and (C) base-Hsm3-Nas6 assembly states, with protein compositions depicted in the inset cartoon. The flexible ATPase domain of Rpt6 is not modeled in (C).

### Nas2 interactions are coupled to completion of the N-ring

With the base-Nas2-Rpn14-Hsm3-Nas6 complex, we were able to trap an intermediate that is normally present only very briefly after association of the Hsm3 module. This proposed first state with all subunits appears very loosely assembled, resulting in limited resolution. Individual modules are held together mainly through chaperone-mediated interactions and an N-ring-like structure, while the ATPase ring is not formed (Fig. 3A and 4A; Supplemental Video 1). The AAA+ motor domains of Rpt6, Rpt3, Rpt4, and Rpt5 are positioned in roughly the right orientation to each other, yet with wide open interfaces. Notably, the ATPase domains of Rpt1 and Rpt2 are turned perpendicular relative to the other Rpts and the expected ATPase-ring plane, creating large gaps to their Rpt neighbors. Association of the Hsm3 module thus seems mediated largely by interactions between the non-ATPase subunits Rpn1 and Rpn2, and the Nas2 chaperone. The base-Nas2-Rpn14-Hsm3-Nas6 complex showed clear density for Nas2’s N-terminal α-helical domain embracing the C-terminal small AAA+ subdomain of Rpt5 (Fig. 4A; Supplemental Video 3). Similar to a previously solved crystal structure of a complex formed by fragments of Nas2 and Rpt5 ^17^, we observed that helices α3 and α4 of Nas2’s N-terminal domain engage in hydrophobic and electrostatic interactions with helices α3 and α4 of Rpt5’s small AAA+ subdomain (Fig. 4B). While the C-terminal tail of Rpt5 with the HbYX motif for 20S core binding remained unresolved in the crystal structure, we were able to use an AlphaFold prediction to model this tail into our cryo-EM density at the chaperone-Rpt5 interface, forming hydrophobic interactions with α3 and α4 of Nas2 (Extended Data Fig. 7A, B).

**Figure 4.**
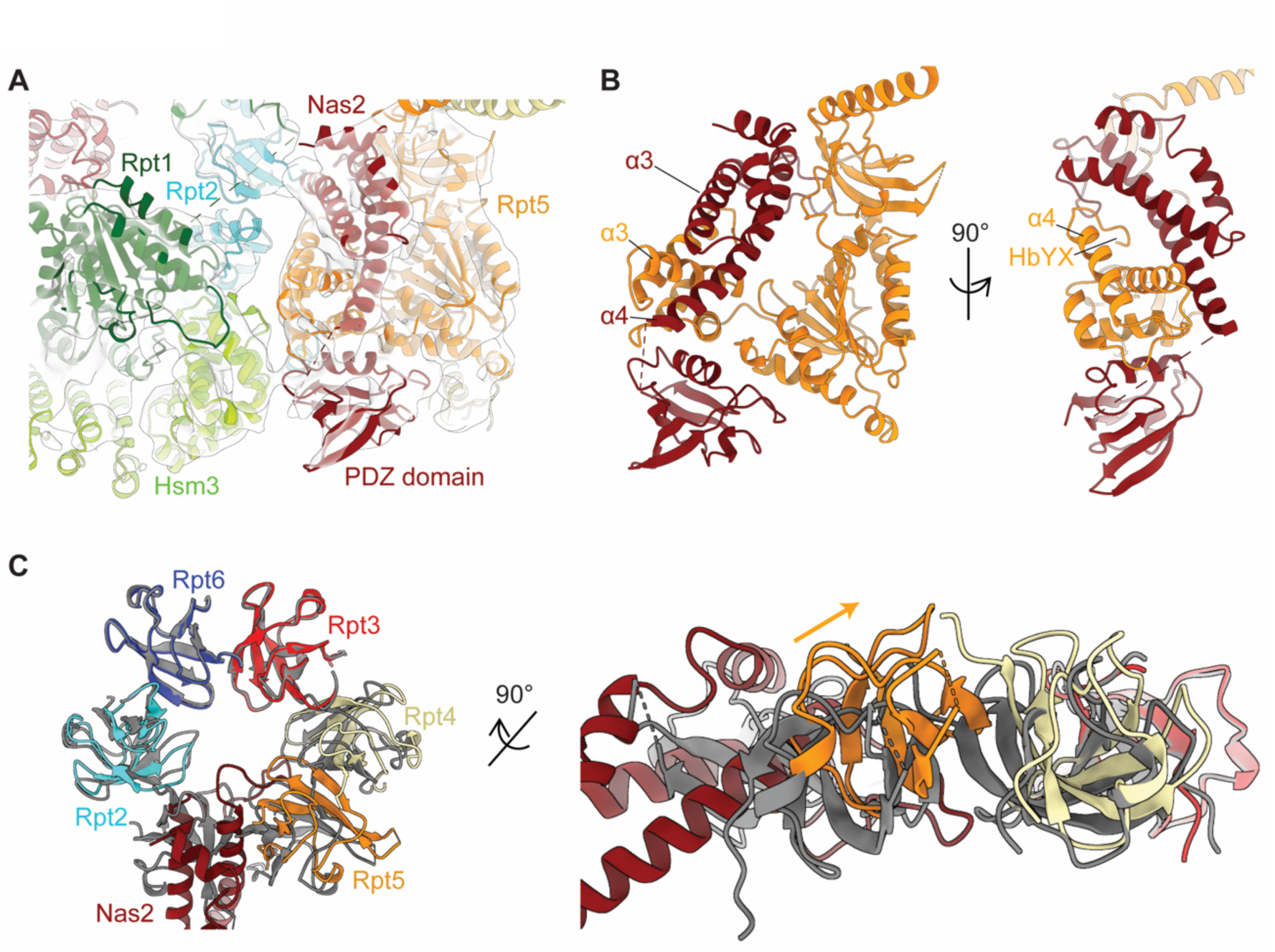
Nas2 interactions with Rpt5 and its participation in the N-ring. (A) Close-up view of Nas2 and Rpt5 in the base-Nas2-Rpn14-Hsm3-Nas6 state (Fig. 3A). Nas2’s N-terminal helices interact with Rpt5’s NTD and small AAA+ subdomain, while Nas2’s PDZ domain binds to both the small and large AAA+ subdomains of Rpt5. The PDZ domain also contacts the neighboring Hsm3 that is in complex with Rpt1 and Rpt2. In the N-ring, Nas2’s N-terminal helices also interact with Rpt2’s NTD and replace the NTD of Rpt1, which is therefore not resolved in this assembly state. (B) Structural model of Nas2-Rpt5 interactions during base assembly. Helices α3 and α4 of Nas2 and helices α3 and α4 of Rpt5 small AAA+ subdomain form the majority of contacts. The HbYX motif at Rpt5’s C-terminus is sandwiched within this interface. (C) Nas2 presence and release are coupled to the maturation of the N-ring. The N-ring of Rpt NTDs in the base-Nas2-Rpn14-Hsm3-Nas6 assembly state (colored) is compared to the N-ring in the base-Rpn14-Hsm3-Nas6 state (grey) after Nas2 was released and Rpt1’s NTD took its spot in the ring. The NTDs of Rpt4 and Rpt5 are tilted upward in the presence of Nas2, but form a planar ring after Nas2 release.

Interestingly, with the extended part of its N-terminal domain with α3, α4, and the loop connecting these helices, Nas2 is part of the N-ring of ATPase subunits, interacting with the NTD’s of Rpt2 and Rpt5 on either side (Extended Data Fig. 7C; Supplemental Video 3), and taking the position that is normally occupied by Rpt1’s NTD in the fully assembled base. Nas2 therefore also prevents an interaction between the NTDs of Rpt1 and Rpt2, which are held together only through contacts of their N-terminal coiled coil and the Hsm3 chaperone bridging their ATPase domains. The participation of Nas2 in the N-ring causes a slight upward tilt for the NTDs of Rpt4 and Rpt5, and a shallow left-handed spiral pitch of the entire N-ring (Fig. 4C). This distortion and the occupancy of Rpt1’s spot in the N-ring may drive the rapid ejection of Nas2 after association of the Hsm3 module, which then allows the formation of the complete N-ring as a rigid hub within the base complex.

On the other end of Nas2’s N-terminal domain, the α4 helix transitions to a less well-resolved density that with the help of an AlphaFold model could be assigned to the C-terminal PDZ domain of Nas2. This PDZ domain interacts with both the large and small AAA+ subdomains of Rpt5 (Fig. 4A; Supplemental Video 3), which leads to an upward tilting of the entire ATPase domain and its pore loop pointing toward the N-ring. In addition, Nas2 appears to form a small interface with the neighboring Rpt1-bound Hsm3 chaperone. Combined with its direct contacts of Rpt5 and Rpt2, this explains why association of the Hsm3 module during base assembly depends on the presence of both the Rpn14/Nas6 and Nas2 modules.

### Rpn14 forms extensive interfaces within the Rpn14/Nas6 module

Both assembly states, base-Nas2-Rpn14-Hsm3-Nas6 and base-Rpn14-Hsm3-Nas6, show density for Rpn14 bound to Rpt6 (Fig. 3A, B). Local refinement with a mask around the Rpt6-Rpn14 portion of the Nas2-bound base resulted in a map for Rpt6-Rpn14 at 3.1 Å resolution, which allowed us to build a molecular model and analyze the interactions of the chaperone with the ATPase subunit (Fig. 5A). The structure of Rpn14 in this complex is essentially identical to a previous crystal structure of Rpn14 alone, showing little flexibility or dynamics within the protein (Extended Data Fig. 8A). However, Rpn14’s interactions with Rpt6 differ substantially from previous models. Those prior studies predicted contacts only between the Rpn14 C-terminal WD40 domain and Rpt6’s small AAA+ subdomain ^30,31^. Instead, we observed extensive interactions between Rpn14’s N-terminal domain and Rpt6’s small AAA+ subdomain, while Rpn14’s C-terminal WD40 domain primarily interacts with the large AAA+ subdomain and forms a minor interface with the small subdomain (Fig. 5A; Supplemental Video 4). Multiple acidic residues in Rpn14’s N-terminal domain and the C-terminal region of the WD40 propeller form a large negatively charged interface that contacts a positively charged surface on Rpt6’s small AAA+ subdomain (Extended Data Fig. 8B), and similarly, another patch of negative charges on the WD40 domain contacts several positively charged residues on Rpt6’s large AAA+ subdomain (Extended Data Fig. 8C). In addition, Met158 in Rpn14 forms hydrophobic interaction with the helical residues 171-179 in Rpt6 (Fig. 4B left), which in the fully assembled ATPase hexamer are part of the interface with Rpt2 (Fig. 4B right). Rpn14 dissociation from Rpt6 is therefore required to eventually close the gap between Rpt2 and Rpt6 and form a functional ATPase ring. Our data also reveal how Rpn14’s Asp157 shapes the loop that contains Met158, which explains previously identified role of Asp157 in Rpt6 binding despite being buried ^30^.

**Figure 5.**
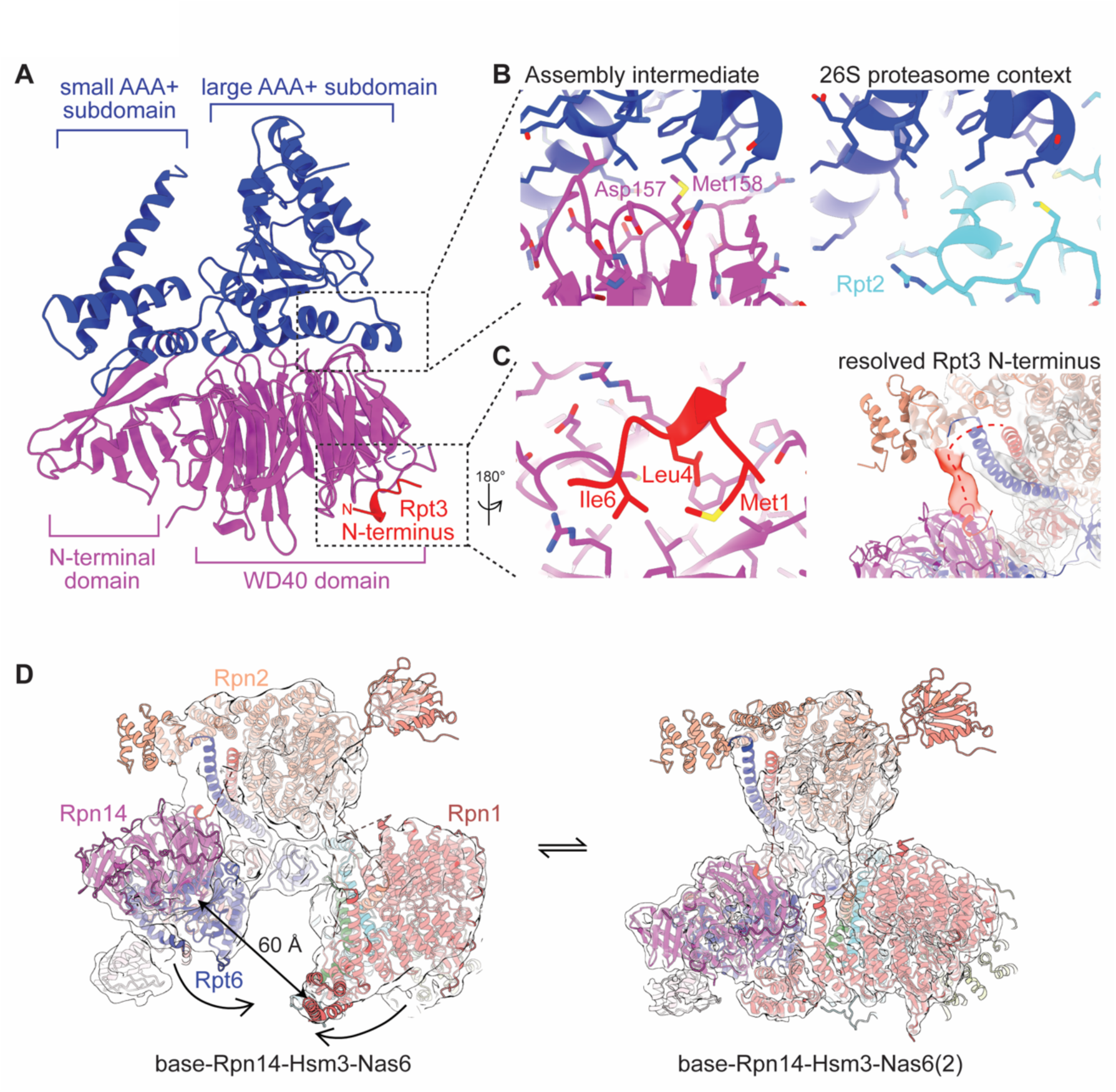
Rpn14 forms extensive interaction within the Rpn14/Nas6 module and binds Rpn1. (A) Cryo-EM based structural model for the complex between Rpn14 (magenta) and Rpt6 (blue). The N-terminal domain of Rpn14 interacts with Rpt6’s small AAA+ subdomain, while its C-terminal WD40 domain mainly binds Rpt6’s large AAA+ subdomain. Rpt3’s N-terminus is shown in red interacting with Rpn14’s WD40 domain. (B) The interactions between Rpn14 and Rpt6 compete with Rpt6 binding to Rpt2. Left: zoom-in view of the interactions between Rpt6 helical residues 171-179 and Rpn14 Met158, with Asp157 stabilizing this conformation. Right: Interactions between Rpt6 and Rpt2 in a mature base within the 26S proteasome (PDB: 6EF1) ^5^. (C) Rpt3’s N-terminus docks on top of Rpn14. Left: zoom-in view of Rpt3 N-terminal residues 1-6 contacting Rpn14’s two propeller blades formed by residues 139-178 and 179-217, with Met1, Leu4, and Ile6 forming hydrophobic interactions. Right: map of base-Rpn14-Hsm3-Nas6 state shows continuous density (red) connecting Rpt3 coiled-coil domain to the interaction site with Rpn14. (D) Large-scale motions of Rpn14-Rpt6 and Rpn1 result in Rpn14-Rpn1 interactions. In the base-Rpn14-Hsm3-Nas6 conformation, there is a ∼ 60 Å gap between Rpn14 and Rpn1’s N-terminus. This gap is closed in the base-Rpn14-Hsm3-Nas6(2) conformation as a result of Rpn14-Rpt6 moving away from Rpn2 and Rpn1 moving away from Rpt1-Rpt2.

Intriguingly, we observed additional density sandwiched between two blades of Rpn14’s WD40 propeller formed by residues 139-178 and 179-217. Based on AlphaFold predictions and subsequent model building, we identified this extra density as the end of Rpt3’s long intrinsically disordered N-terminus, forming hydrophobic interaction on top of Rpn14 (Fig. 4C left). Consistently, we also observed a continuous connecting density from the Rpt6/Rpt3 coiled-coil to the binding site on top of Rpn14 (Fig. 4C right).

Overall, our cryo-EM structures revealed that Rpn14’s interactions within the Rpn14/Nas6 module are much more extensive than previously assumed. Rpn14 covers a large portion of the outward-facing surface on Rpt6’s large and small AAA+ subdomains, and thus rigidifies the conformation of this ATPase subunit. Moreover, the interactions of Rpn14 with the N-terminus of Rpt3 may provide an additional mean of proofreading for the correct assembly of the Rpn14/Nas6 module.

Further classification of particles also revealed a second conformation, base-Rpn14-Hsm3-Nas6(2), in which Rpn14-Rpt6 moved away from Rpn2’s N-terminus and Rpn1 is repositioned relative to Rpt1/Rpt2. Importantly, in this state Rpn14 directly interacts with Rpn1, which closes a ∼ 60 Å gap present between those proteins in base-Rpn14-Hsm3-Nas6 (Fig. 5D; Supplemental Video 5). The interface involves the blade residues 181-287 of Rpn14’s WD40 propeller and the helical residues 1-143 at the N-terminus of Rpn1, forming mostly hydrophilic interactions. Compared to the base-Rpn14-Hsm3-Nas6 conformation, Rpt6 and Rpt3 in base-Rpn14-Hsm3-Nas6(2) swing up relative to Rpt4 and Rpt5 to form an overall more planar arrangement for these four ATPase subunits (Extended Data Fig. 9A). On the other hand, Rpt1 and Rpt2, together with Hsm3, become much more dynamic, as Rpn1 swings away to interact with Rpn14 (Extended Data Fig. 9B). Rpn14 may hence couple the movement of Rpn1 to the movement of ATPase domains in the open AAA+ ring of the base-Rpn14-Hsm3-Nas6 state, potentially facilitating their transition to a closed motor conformation in the base-Hsm3-Nas6 state.

### Hsm3 cradles Rpt1 and Rpt2

To understand how Hsm3 is chaperoning Rpt1 and Rpt2 during base assembly, we locally refined particles of the base-Hsm3-Nas6 complex with a mask around Hsm3 and the ATPase domains of Rpt1 and Rpt2, and achieved a nominal resolution of ∼ 3.0 Å (Extended Data Fig. 10A). Local refinement using a similar mask on the base-Rpn14-Hsm3-Nas6 complex returned essentially the same density (Extended Data Fig. 10B, C), indicating that Rpt1, Rpt2, and Hsm3 move as a rigid body from the open ATPase conformation in base-Rpn14-Hsm3-Nas6 to the more closed conformation in base-Hsm3-Nas6. In agreement with the previously solved crystal structure of Hsm3 in isolation ^33^, the chaperone adopts a horseshoe-shaped structure in the module or base context as well. With its 11 HEAT repeats Hsm3 wraps around the large and small AAA+ subdomains of Rpt1, while also interacting with the large and small AAA+ subdomains of Rpt2 through the outside of its three C-terminal HEAT repeats (Fig. 6A, B; Supplemental Video 6). Hsm3 thereby uses its N-terminal helices including residues 1-86 to form mixed hydrophilic and hydrophobic interactions with Rpt1 helix 214-239 in the large AAA+ subdomain (Extended Data Fig. 11A). In addition, a hydrophobic patch on the last two helices of Hsm3 contacts Phe319 in Rpt1’s pore-2 loop (Extended Data Fig. 11D). The interactions with Rpt1’s small AAA+ subdomain largely agree with a previous crystal structure for Hsm3 in complex with a C-terminal fragment of Rpt1 ^32,33^, whereby a hydrophobic core is surrounded by electrostatic interactions (Extended Data Fig. 11B, C).

**Figure 6.**
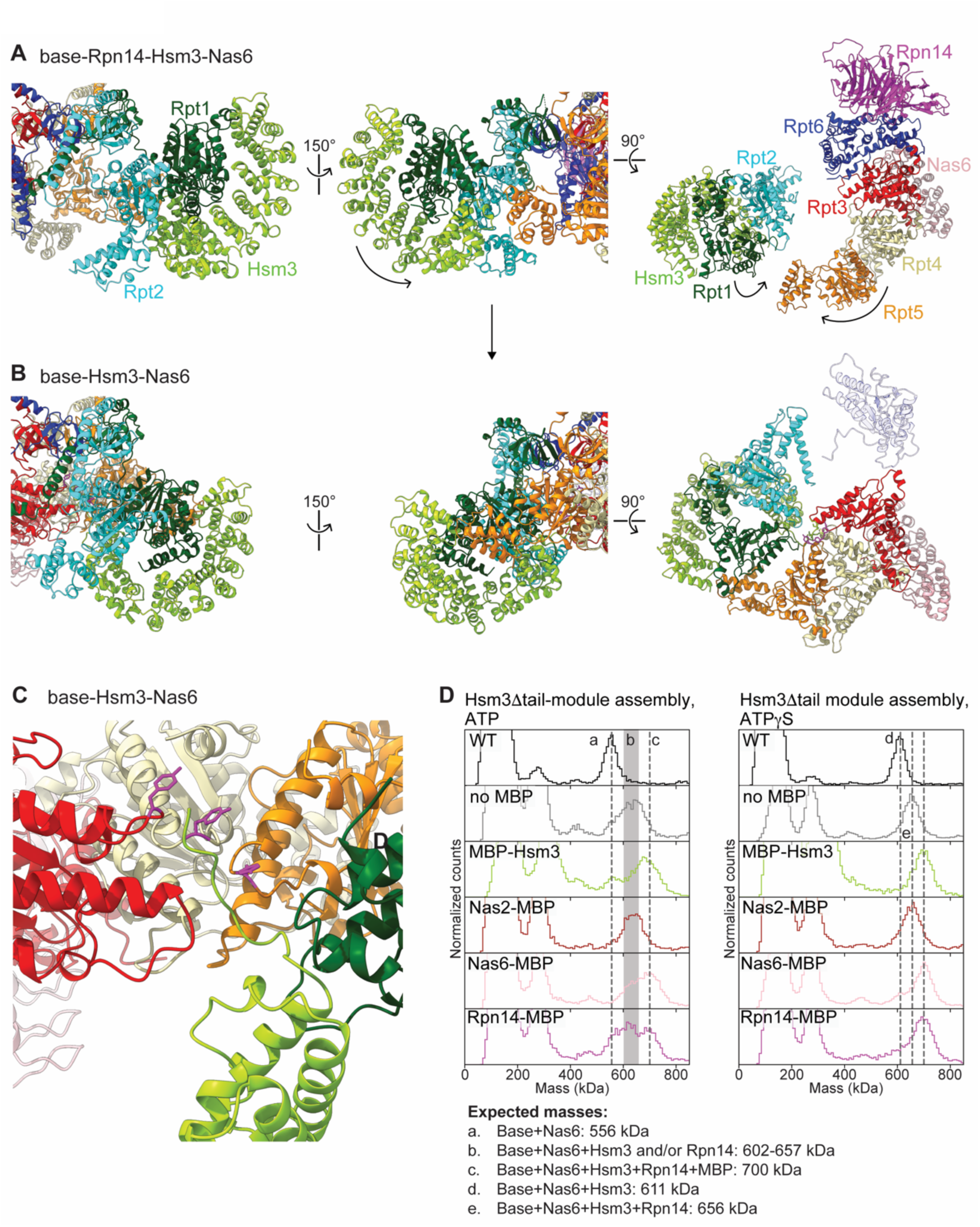
Hsm3 guides closing of the AAA ring and allosterically activates ATPase activity through its C-terminal tail. (A, B) Comparison of Hsm3-module conformations between (A) the base-Rpn14-Hsm3-Nas6 state and (B) the base-Hsm3-Nas6 state. Left: views from the Rpn1 side (with Rpn1 removed) show pivoting of the Hsm3-bound ATPase domains of Rpt1 and Rpt2 as a rigid body from a perpendicular position to being more planar with the other ATPase domains. Middle: view from Rpt1-Rpt5 side. The same rotation as in the left panel, highlighting the closing of the gap between Rpt1 and Rpt5 as a result. Right: view from the top of the AAA ring. Only the ATPase domains and chaperones are depicted. The ATPase domains of Rpt3, Rpt4, and Rpt5 move to close their gap to the neighboring subunits, while Rpt1 and Rpt2 rotate to be in the same plane, resulting in a conformation resembling mature base ATPase ring. Rpt6 (shown transparent) becomes highly dynamic and cannot be confidently modeled in the base-Hsm3-Nas6 state. (B) Hsm3’s C-terminal tail mimics a substrate polypeptide in the central channel, interacts with the pore-1 loop tyrosines (purple) of Rpt3, Rp4, and Rpt5, and arranges them into a spiral staircase that resembles a substrate-processing state. (D) Hsm3’s C-terminal tail is important for activating the ATPase activity of the base and allosterically releasing Rpn14. The importance of Hsm3’s tail on chaperone release was analyzed by MBP mass shift assay as in Fig. 1C, with the assembly containing wild-type Hsm3 as control (WT) and all other assembly reactions performed with Hsm3Δtail. Where indicated, MBP was attached to the specified chaperone. Left: assembly in the presence of ATP shows that Hsm3’s tail is important for the release of both Hsm3 and Rpn14. The shaded area b includes several species that have similar masses and cannot be distinguished. Right: assembly in the presence of ATPγS shows that Rpn14 release is compromised by the lack of Hsm3’s tail independent of ATP hydrolysis.

**Figure 7.**
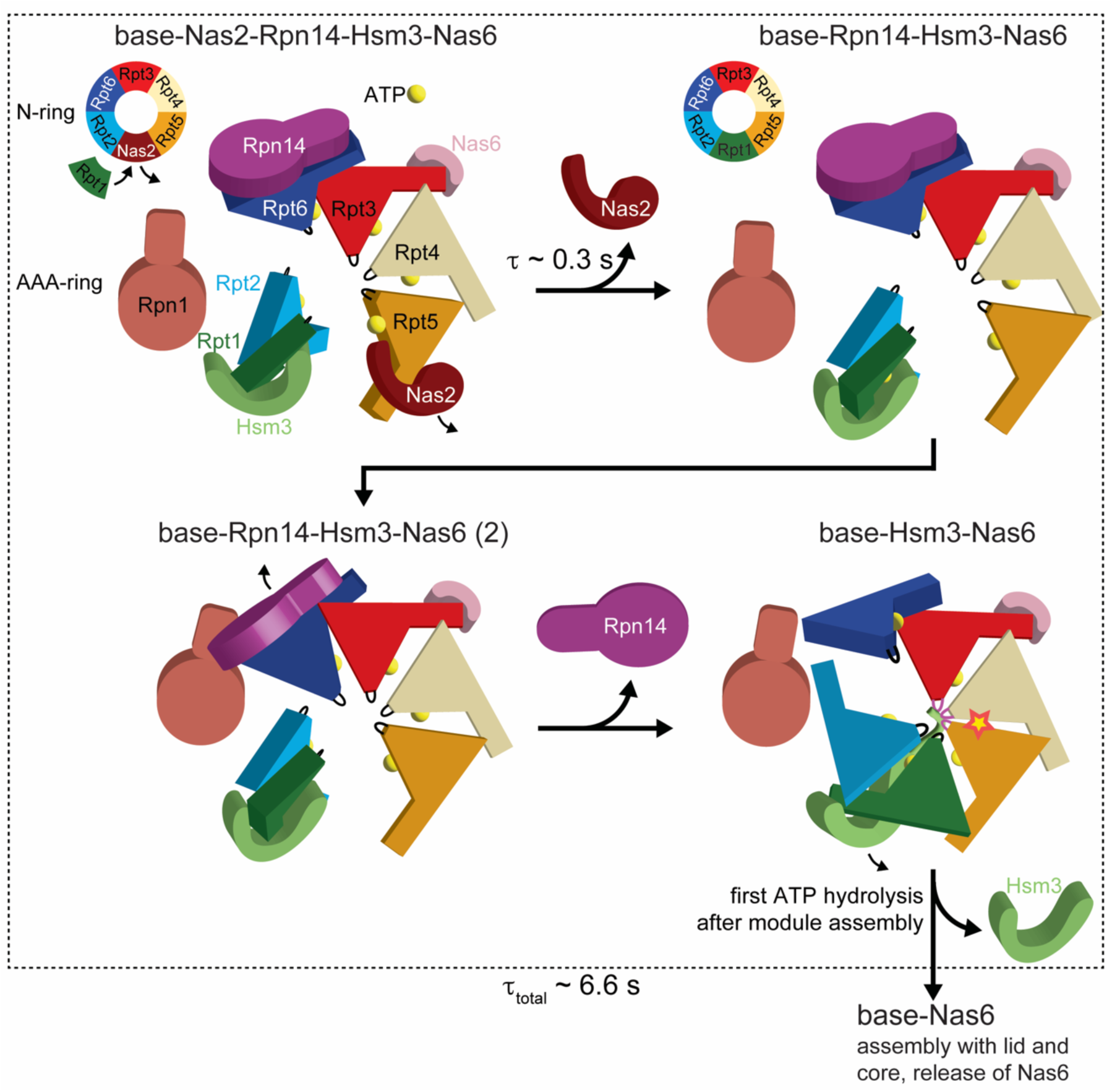
Schematic model for the assembly of the 26S proteasome base. After association of the Hsm3 module with the Nas2 and Rpn14/Nas6 modules, the AAA-ring is only loosely assembled and the Rpt subunits are primarily held together by the N-ring, in which the N-terminal domain of Nas2 occupies the space of Rpt1’s N-domain (top left). All six Rpts are occupied with ATP (yellow spheres). Nas2 is then quickly released, leading to completion of the N-ring and an even more open AAA-ring, with the ATPase domains of the Hsm3-bound Rpt1 and Rpt2 oriented perpendicular to the other 4 subunits (top right). Subsequent interactions between Rpn14 and Rpn1 partially close the large gap between Rpt6 and Rpt2 (bottom left), before the C-terminal tail of Hsm3 induces a more closed AAA-ring conformation that resembles a substrate-processing state and ejects Rpn14 (bottom right). In this state, Hsm3’s tail mimics a substrate polypeptide in the central channel and interacts with the pore-1 loops (purple) of Rpt3, Rpt4, and Rpt5, which are arranged in a righthanded spiral staircase. Only Rpt4 has a closed ATPase site that is competent for hydrolysis (red star), and the conformational changes induced by this hydrolysis release Hsm3 ∼ 6.6 s after the Hsm3 module associated with the other two modules. The resulting Nas6-bound base is active in ATP hydrolysis and competent to bind the core and lid subcomplex for formation of the 26S proteasome holoenzyme.

Hsm3’s contacts with the bottom of Rpt2’s ATPase domain are consistent with a previous domain-deletion study ^32^ and include hydrophobic interactions, potential hydrogen bonding, and electrostatic interactions (Extended Data Fig. 11E). As a result, Hsm3 bridges Rpt1 and Rpt2, while preventing their ATPase domains from extensive direct contacts and holding them in a stacked rather than adjacent orientation. The minimal direct interactions between Rpt1 and Rpt2 involve only their N-terminal coiled coil and a small interface between Rpt1’s small AAA+ subdomain and Rpt2’s helix 204-212. In addition to keeping Rpt1 and Rpt2 apart, Hsm3 also prevents Rpt1 docking against Rpt5, which in the closed ATPase hexamer contacts the same Rpt1 helix around residues 214-239 that Hsm3 is occupying.

### Hsm3 C-terminal tail mimics substrate for ATPase-ring closure and activation

The different states observed in our data set provide insights into how the presence of chaperones and their ordered release direct the conformational changes toward a closed, ATP-hydrolysis-active motor that is able to bind lid and core. Comparing the structures for base-Nas2-Rpn14-Hsm3-Nas6 and base-Rpn14-Hsm3-Nas6 indicates that the eviction of Nas2 allows the insertion of Rpt1’s N-terminal domain into the N-ring, yet further opens up the loosely assembled ATPase ring by eliminating the interaction between Nas2 and Hsm3 (Fig. 3A, B; Supplemental Video 7). The conformational differences between base-Rpn14-Hsm3-Nas6 and base-Rpn14-Hsm3-Nas6(2) suggest that the direct contact subsequently established between the Rpt6-bound Rpn14 and Rpn1 pulls the Rpn14/Nas6 and Hsm3 modules closer together and potentially prepares the ATPase ring for closure (Supplemental Video 5). The following transition to base-Hsm3-Nas6 not only involves the release of Rpn14, but is also linked with a rotation of the ATPase domains for Rpt1 and Rpt2 (Fig. 6A, B, left and middle panels; Supplemental Videos 8 and 9), from a perpendicular orientation to lying within the ring plane, and with a partial closure of the gaps between Rpt3, Rpt4, and Rpt5 (Fig. 6A, B, right). Interestingly, local refinement of the base-Hsm3-Nas6 state using a mask on the six ATPase domains revealed that Hsm3’s C-terminal tail occupies the bottom section of the central channel, which is also in agreement with recent *in vivo* crosslinking results ^34^. The tail is thereby tightly engaged by the pore-1 loop tyrosines of Rpt3, Rpt4, and Rpt5 in a spiral-staircase arrangement (Fig. 6C; Supplemental Video 10), while Rpt1 and Rpt2 are unengaged toward the bottom of the staircase, and Rpt6 is highly mobile between Rpn1 and Rpt3. Hsm3’s tail thus appears to mimic a substrate polypeptide and induce an Rpt arrangement that partially resembles a substrate-processing motor conformation, with several subunits in a staircase and Rpt6 as the mobile seam subunit on its way to the top. In this state, Phe315 of Rpt4 and Phe315 of Rpt5 are flipped to engage with the neighboring Rpt3 and Rpt4, respectively, which also resembles the Rpt interactions observed during spiral-staircase movements in the substrate-processing proteosome ^5^. Hsm3’s C-terminal tail thus seems to coordinate the conformational transitions of ATPase domains from an open ring in base-Rpn14-Hsm3-Nas6 to an almost closed ring in base-Hsm3-Nas6 that is accompanied by the release of Rpn14.

To confirm the importance of the C-terminal tail for facilitating these transitions, we created a Hsm3 module with the last 16 residues deleted from Hsm3 (Hsm3Δtail) and analyzed its assembly with the other modules by again using MBP fusions of individual chaperones in mass photometry experiments. Indeed, the lack of Hsm3’s C-terminal tail caused a mass shift corresponding to Hsm3Δtail’s molecular weight for all assembled base complexes and an additional MBP-equivalent shift for the assembly with the MBP-Hsm3Δtail construct (Fig. 6D), confirming the retention of tailless Hsm3. Intriguingly, Rpn14 was also retained to a considerable extent on the base assembled with Hsm3Δtail in ATP (Fig. 6D left), and stayed even quantitatively bound when performing this assembly in the presence of ATPγS (Fig. 6D right).

In the base-Hsm3-Nas6 state, the Rpt2/Rpt6 and Rpt6/Rpt3 ATPase interfaces are not formed due to Rpt6’s high mobility, and the interfaces of Rpt3/Rpt4, Rpt5/Rpt1, and Rpt1/Rpt2 are also still open to variable extent (Extended Data Fig. 12A-C, E). Rpt5, Rpt1, Rpt2, Rpt6, and Rpt3 are therefore likely incompetent to hydrolyze ATP. In contrast, the interface of Rpt4/Rpt5 is closed and has all interactions established around the ATP-binding site to allow hydrolysis (Extended Data Fig. 12D). Together with our finding that ATP hydrolysis in Rpt4 is required to eject Hsm3 and complete the base assembly process, this indicates that Rpt4 is the first ATPase subunit to hydrolyze ATP, which likely rearranges the clockwise-neighboring Rpt5 and Rpt1 in the staircase, closes their interfaces, and displaces Hsm3. By using its C-terminal tail to mimic a substrate polypeptide and induce a processing conformation of the ATPase ring, Hsm3 may thus promote the ATP hydrolysis necessary for its own release and for maturation of the base. Consistent with a role of its C-terminal tail in allosterically activating the ATPase ring, we observed a ∼ 30% lower ATPase activity for base that was assembled with Hsm3Δtail and consequently retained this chaperone (k_ATPase_^Δtail^ = 0.71 +/- 0.04 s^-1^ vs. k_ATPase_^WT^ = 1.06 +/- 0.01 s^-1^; Extended Data Fig. 1B). Combined with our mass-photometry measurements, these results suggest a model where the processing state-like base conformation induced by the Hsm3 tail promotes efficient Rpn14 release independent of ATP hydrolysis, followed by the ejection of Hsm3 that requires ATPase activity in Rpt4. In the absence of Hsm3’s tail, the base adopts an unproductive state that only partially expels Rpn14 upon ATP hydrolysis, yet completely fails to release Hsm3. In summary, the C-terminal tail of Hsm3 thus plays critical roles in inducing a closed Rpt-ring conformation with hydrolysis-competent interfaces, which allosterically promotes the ejection of Rpn14 and allows the subsequent ATP-hydrolysis-driven release of Hsm3.

## Discussion

Assembly of the proteasomal heterohexameric AAA+ motor presents a unique conundrum: while interactions between the Rpts need to be stable and specific to arrange the subunits in a defined order, a large portion of their interfaces have to stay dynamic to allow for ATP-hydrolysis-driven motor movements during substrate processing. Static interfaces are limited to the N-terminal coiled coils that determine the formation of specific Rpt dimers and to the OB-fold-containing N-domains, which contribute only ∼ 600 A#^2^ per subunit interface. The eukaryotic cell adopts a modular solution to this assembly challenge by dividing the six subunits into three pairs of dimers, each with dedicated chaperones to stabilize the modules themselves and regulate their correct association, in part by forming inter-module contacts. This strategy transfers the complexity from fine-tuning the strength of subunit interactions to ensuring chaperone release at the right stage of assembly. Indeed, overexpression or addition of Nas6, Rpn14, or Hsm3 inhibits 26S proteasome activity ^39,40^, suggesting an intricate balance between module stability, assembly, and timely chaperone release to prevent interference with holoenzyme formation. Our data point to a model where chaperone interactions stabilize subunit conformations and orientations that are strongly distinct from those in the mature base subcomplex. Upon association of all three modules, the chaperones guide the intermediates across the conformational landscape toward the mature state, with chaperone release at key checkpoints along the way. The large conformational changes thereby allow the chaperones to bind the modules and assembly intermediates with high affinity, while only weakly interacting with the mature base. Interestingly, in our structures we observed a limited number of globally distinct conformations for each state of the assembly process, despite the large conformational space that is theoretically accessible to the loosely associated modules. The presence of chaperones thus appears to restrict the conformational space sampled by the intermediates, reduce unproductive states, and funnel the dynamics toward a specific pathway. Each release of a chaperone may open up a new conformational space to be sampled, leading to sequential, coordinated chaperone dissociations. This is in sharp contrast to the previously prevailing assumption that chaperones bind intermediates that largely resemble the conformation of the mature base subcomplex ^15,21,26,39–41^.

The most extreme chaperone-regulated conformational changes occur during rearrangements of the AAA+ ATPase ring. During the transition from the base-Nas2-Rpn14-Hsm3-Nas6 state to the base-Rpn14-Hsm3-Nas6 and base-Rpn14-Hsm3-Nas6(2) conformations, and then to the base-Hsm3-Nas6 state, the AAA+ domains gradually establish inter-subunit contacts that resemble the mature base, shedding chaperones in the process (Supplemental Video 11). In the Nas2- and Rpn14-bound states, none of the Rpt interfaces are closed and ATPase sites lack the arginine fingers from the neighboring subunit required for hydrolysis. This structural feature also explains why individual modules are inactive in ATP hydrolysis, which prevents the waste of energy in the cell. The ATPase-domain separation between neighboring subunits is thereby structurally enforced by the chaperones. Hsm3 binding requires Rpt1 and Rpt2 to be apart, with Rpt1 encircled by Hsm3’s HEAT repeats, while Rpt2 docks against the side. Nas2 binds the N-domain and both AAA+ subdomains of Rpt5, tilting up the entire ATPase domain and preventing Rpt4 from engaging. Rpn14’s effects are more subtle: Rpn14 has an extensive interface with both the small and large AAA+ subdomains of Rpt6 and likely biases Rpt6’s conformation against forming the ATPase interface with Rpt3. Furthermore, Rpn14 binds the N-terminus of Rpt3 and thus establishes a connection between Rpt6’s ATPase domain and the Rpt6/Rpt3 coiled coil through Rpt3’s N-terminal IDR, which restricts the conformational freedom of Rpt6. This limited motion likely keeps Rpt6’s pore-loop pointing downward and prevents the AAA+ domain from engaging with Rpt3. In contrast, Nas6 interaction with the C-terminal small AAA+ subdomain of Rpt3 is compatible with a wide range of conformations and does probably not contribute to holding Rpt3 apart from Rpt6 or Rpt4. Instead, the Rpt3-Rpt4 separation may result from the concerted effects of Nas2 and Rpn14 on Rpt5 and Rpt6, respectively.

Interestingly, while Nas2, Nas6, and Hsm3 are conserved throughout eukaryotes, no ortholog of Rpn14 is found in plants ^42^ and some species of fungi and animals, suggesting a recent evolution of Rpn14 in the common ancestor of animals and fungi, and an occasional loss of Rpn14 thereafter. The lack of Rpn14 is also accompanied by a loss in conservation of Rpt3’s N-terminus (Extended Date Fig. 13), which is in agreement with the direct interaction we observed for the two proteins. How species without Rpn14 enforce the separation of Rpt6 and Rpt3 during base assembly remains unclear, but it is possible that this role is taken on by Nas6 or that the Rpt6/Rpt3 interface is intrinsically weaker in these species.

Macromolecular assembly processes are often driven by chemical reactions, like nucleotide hydrolysis, that occur directly in the macromolecule or in the involved chaperones to provide the energy for conformational changes or the switching between different affinity states. The base subcomplex utilizes its own AAA+ ATPase activity to promote assembly and chaperone ejection, with Hsm3 as the major regulator. Hsm3 release requires ATP hydrolysis, and Hsm3’s C-terminal tail functions as an allosteric activator of this ATPase activity by mimicking a substrate and arranging Rpt3, Rpt4, and Rpt5 into a processing conformation. Intriguingly, this is reminiscent of other macromolecular assembly chaperones. For example, formation of the large ribosomal subunit involves multiple chaperones that help shaping the peptide-exit tunnel by inserting themselves into the growing tunnel and mimicking a nascent polypeptide, before getting extracted by assembly motors during maturation ^43,44^. These observations suggest a common principle, where chaperones simultaneously template the assembly and test the quality of the resulting product by mimicking a substrate of the macromolecule. In case of the proteasomal base subcomplex, Hsm3 appears to conduct a “dry run” that tests the complex’s ability to adopt a substrate-processing state with a spiral staircase around Hsm3’s tail and to hydrolyze ATP in order to release this tail together with the entire chaperone. If this quality control fails, Hsm3 remains bound and sterically interferes with holoenzyme assembly, such that dysfunctional base subcomplexes are prevented from poisoning the 26S proteasome pool. Since the dissociation of chaperones is directly coupled to the successful assembly and maturation of the proteasome base, long-lived chaperone-bound intermediates may be recognized as defective and targeted for degradation. This would be consistent with recent studies showing that macromolecular assembly processes are intrinsically error-prone and require surveillance by quality-control pathways ^45,46^. Indeed, mammalian Rpn14 was found to be targeted by the UPS for the turnover of orphaned Rpt6-Rpt3 ^47^. It is possible that Nas2 and Hsm3 also serve as markers for quality control, but the exact selection criteria, the proteins involved, and the eventual fate of faulty assemblies remain to be investigated.

## Methods

### Cloning of base module components

The plasmids used for the recombinant expression of the base-assembly modules were similar to those previously described^37^. Briefly, yeast Rpn1, Rpn2 and Rpn13 were cloned into a pETDuet-1 plasmid, and Nas2, Nas6, Hsm3 and Rpn14 were cloned into a pACYCDuet-1 plamid. A T7 promoter was cloned before each subunit, and one T7 terminator was cloned at the end of the multiple cloning sites. The pACYCDuet-1 also contained genes encoding rare tRNAs to account for differences in codon usage between yeast and *E. coli*. Genes encoding yeast Rpt1/Rpt2, Rpt6/Rpt3 and Rpt4/Rpt5 were separately cloned into a pCOLA-1 vector, with each Rpt gene having its own T7 promotor and terminator. A His_6_ tag was added to the N-terminus of Rpt1, Rpt3, and Rpt4, while a FLAG tag was added to the N-terminus of Rpt2, Rpt6, and Rpt5. For experiments involving MBP-tagged chaperones, the sequence coding for *E. coli* MBP was attached to the N-terminus of Hsm3 or the C-terminus of Nas2, Nas6 and Rpn14, and a short flexible Gly/Ser linker was placed between MBP and the chaperones. For fluorophore labeling of Nas2 and Hsm3, the ybbr13 tag sequence was attached to the C-terminus of Nas2 or the N-terminus of Hsm3. For Walker-B mutation in the Rpt subunits, a glutamate to glutamine substitution was introduced at residues 310, 282, 249, 273, 282 and 282 in Rpt1, Rpt2, Rpt6, Rpt3, Rpt4, and Rpt5, respectively.

### Expression and purification of assembly modules

A previous protocol for the preparation of the recombinant base subcomplex was slightly modified for the preparation of modules ^37^. Briefly, *E. coli* BL21(DE3)* was co-transformed with the plasmids coding for Rpn1, Rpn2, Rpn13, Nas2, Nas6, Rpn14 and Hsm3, and electrocompetent cells were prepared from the resulting clone. Those cells were further transformed with plasmids coding for either Rpt1/Rpt2, Rpt6/Rpt3, or Rpt4/Rpt5 to allow expression of the Hsm3 module, Rpn14/Nas6 module, or Nas2 module, respectively. The transformed cells were grown in 4 L of dYT medium at 37 ℃ to an OD_600_ = 0.5. Expression was induced with 1 mM IPTG, and cells were incubated at 30 ℃ for 5 hours and then overnight at 16 ℃. Cells were pelleted and resuspended in NiA buffer (60 mM HEPES, pH 7.6, 100 mM NaCl, 100 mM KCl, 10 mM MgCl_2_, 5% glycerol, 1 mM ATP, and 20 mM imidazole) supplemented with benzonase and protease inhibitors (aprotinin, pepstatin, leupeptin, and AEBSF). After lysis by sonication, the lysate was centrifugated for 30 min at 30,000 g. The supernatant was then loaded on a 5 mL HisTrap FF crude column and washed with 50 mL NiA buffer, before elution with 15 mL NiA buffer supplemented with 250 mM imidazole. The eluent was then loaded on a 5 mL column with anti-Flag M2 affinity resin, washed with 50 mL NiA buffer, and bound proteins were eluted with 15 mL NiA buffer supplemented with 0.3 mg/mL 3×FLAG peptide. After concentrating FLAG eluent to a volume of ∼ 0.5 mL, the sample was loaded on a size exclusion Superose 6 increase 10/300 column equilibrated in GF buffer (30 mM HEPES, pH 7.6, 50 mM NaCl, 50 mM KCl, 10 mM MgCl_2_, 5% glycerol, 0.5 mM TCEP, and 1 mM ATP) and eluted in the same buffer. The size exclusion fractions corresponding to modules were collected and concentrated. The concentrations of individual modules were determined by Bradford assay using bovine serum albumin (BSA) as a standard. All modules, including the ones with modifications, were purified by the same protocol.

### Fluorophore labeling and biotin modification

Fluorophores were attached to Hsm3 and Nas2 by Sfp labeling as previously described ^48^. Coenzyme A (CoA) was incubated with an excess of maleimide-LD555 (a sulfo-Cy3 derivative from Lumidyne Technologies) or maleimide-LD655 (a sulfo-Cy5 derivative from Lumidyne Technologies), before quenching the excess dye with DTT. CoA-dye (200 µM), Sfp (200 µM), and ybbr-tagged modules (50 µM) were mixed in GF buffer for labeling overnight at 4 ℃. The labeled modules were separated from the reaction components using a Superose 6 increase 10/300 column in GF buffer as described for their purification above.

The biotin modification of the Nas2 module was performed using an established protocol^49^. Briefly, the Avi-tag sequence was attached after the His_6_-affinity tag at the N-terminus of Rpt4. Following the Ni-NTA-affinity purification, the Nas2 module was incubated with 200 µM BirA and 1 mM biotin in GF buffer at 4 ℃ overnight (ATP required for the biotinylation reaction was already present in the GF buffer). The biotin-modified module was then further purified by size-exclusion chromatography as described above.

### ATP-hydrolysis measurement

ATPase-hydrolysis activities were measured using an NADH-coupled assay as described previously ^37^. The base subcomplex or individual modules were prepared at 2x concentration of 700 nM in GF buffer and rapidly combined with an equal volume of 2x ATPase mix (1 mM NADH, 7.5 mM phosphoenolpyruvate, 3 U/ml pyruvate kinase and 3 U/ml lactase dehydrogenase), resulting in final concentration of 350 nM for the base or modules. For measurement of module mixtures, one of the modules was kept at a final concentration of 350 nM, with other modules in excess at 500 nM. The samples were transferred to a 384-well clear bottom plate (Corning) and the steady-state depletion of NADH was assessed at 30 ℃ by measuring the absorbance at 340 nm in a Synergy Neo2 Multi-Mode Plate Reader (Biotek).

### Mass photometry

Mass photometry measurements were all conducted on a Refeyn twoMP system using the “Regular” FOV (10.9 x 4.3 µm and 498 Hz) in the AcquireMP software. Sample wells were built by attaching a silicon gasket to the surface of a water/ethanol cleaned glass coverslip.

The experiments were carried out using the droplet-dilution method: A droplet of 9 µL GF buffer without ATP was pipetted into the sample well and focused using the autofocus function in AcquireMP. Then, 1 µL of sample was pipetted into the droplet and mixed, before the movie recording for 60 s was started immediately after the mixing. For measurements in Figure 1A, modules were mixed at 2 µM each in GF buffer and incubated at room temperature for 5 min. The mixtures were quickly diluted to 50 nM in GF buffer and then another 10-fold to 5 nM in GF buffer without ATP on the coverslip (presence of ATP for the final dilution caused high background signal at low molecular weight). For all other measurements, modules were first mixed at 500 nM instead of 2 µM each in GF buffer. For measurement of modules assembled in ATPγS, the modules were mixed and diluted in GF buffer with 1 mM ATPγS, except for the final 10-fold dilution on the coverslip, which was with nucleotide-free GF buffer. The recorded movies were analyzed using DiscoverMP software.

### Cryo-EM grid preparation and data collection

Wild-type (WT) Hsm3 module, WT Rpn14/Nas6 module, and Nas2 module with Rpt4 Walker-B mutation were mixed in GF buffer at 10 µM for each module and incubated on ice for 1 hour. The assembled intermediate was purified from unassembled modules using a Superose 6 increase 10/300 column in GF buffer at 4 ℃. The fractions corresponding to assembled intermediate were pooled and concentrated to ∼ 6 mg/mL based on Bradford assay. Just before grid preparation, 0.01% NP-40 was added to the assembled intermediate solution. UltraAu-foil R1.2/1.3 300-mesh grids (EM Sciences) were glow-discharged at 25 mA for 25 s. Three µL of the assembled intermediate solution was applied to the grid in Vitrobot (Thermo Fisher) maintaining 100% humidity and 4 ℃, waited for 2 s, and blotted once for 3 s using blot force setting 5 before plunge-frozen in liquid ethane.

Grids were clipped and transferred to a Titan Krios G3i operated at 300 kV (Thermo Fisher) equipped with a Gatan K3 camera. Images were taken in SerielEM at a nominal magnification of ×81,000 (1.048 Å pixel size) in super-resolution mode with a defocus ranging from −0.5 to −2.0 μm. We collected 50 frames per shot with a total electron dose of ∼50 e per Å^2^ per s. A total of 8,200 movies were collected for the stalled base assembly intermediate.

### Cryo-EM data processing and model building

Data processing was done in CryoSparc v5.0 (Extended Data Fig. 3). movies were patch-motion corrected and patch CTF estimated. Initially, ∼160,000 particles were picked using a 180-270 Å diameter blob picker. Particles were extracted at 1024 super-resolution pixels and binned to 512 pixels. After 2D classification, ab-initio reconstruction, and heterogeneous refinement, classes showing features and sizes expected for the base assembly intermediates were chosen for template generation. Around 2,000,000 particles were picked using template picker and extracted using a 1024 pixels box, binned to 512 pixels. After multiple rounds of 2D classification, ab-initio reconstruction, and heterogeneous refinement, ∼1,000,000 particles were selected for further refinement. Based on heterogeneous refinement, particles were classified into four classes: base-Nas2-Rpn14-Hsm3-Nas6, base-Rpn14-Hsm3-Nas6, base-Rpn14-Hsm3-Nas6(2), and base-Hsm3-Nas6. Final density maps for each class were generated by doing non-uniform refinement jobs with C_1_ symmetry, using the map and particle class outputs from previous heterogeneous refinements as initial input.

For local maps of Rpn14-Rpt6, Hsm3-Rpt1-Rpt2, or Hsm3-Rpt1-Rpt2-Rpt3-Rpt4-Rpt5, we generated generous masks around the chaperone and ATPase domains of the indicated units using the non-uniform refined density maps, low-passed to 8 Å, and did local refinement jobs using the masks and their corresponding particles. The Rpn14-Rpt6 local refinement was done on maps and particles from the base-Rpn14-Hsm3-Nas6(2) state. We observed that the Rpn14-Rpn1 contact in this state seems to keep Rpn14-Rpt6 less mobile, allowing much better refinement result than for base-Rpn14-Hsm3-Nas6 state. For Hsm3-Rpt1-Rpt2 local refinements, two strategies were pursued: First, maps and particles from the base-Hsm3-Nas6 state non-uniform refinement were used. Second, particles from both the base-Rpn14-Hsm3-Nas6 and base-Rpn14-Hsm3-Nas6(2) states were combined and non-uniform refined with base-Rpn14-Hsm3-Nas6 as the initial structure. This resulted in a density similar to the base-Rpn14-Hsm3-Nas6 state. The pooled particles and their non-uniform refined map were masked for Hsm3-Rpt1-Rpt2 and locally refined. The local map of Hsm3-Rpt1-Rpt2-Rpt3-Rpt4-Rpt5 was obtained from the map and particles of the base-Hsm3-Nas6 state.

All maps directly from CryoSparc refinement jobs without sharpening or enhancement were used for further model building and presentation in the figures. In general, initial models were generated using AlphaFold through its online server ^50^. For the base-Nas2-Rpn14-Hsm3-Nas6 state, we separated the AlphaFold model into three parts: 1. Nas2 and Rpt5; 2. Rpt4, Rpt3, Rpt6, Nas6, Rpn14, Rpn2, and Rpn13; and 3. Rpt1, Rpt2, Hsm3, and Rpn1. For the base-Rpn14-Hsm3-Nas6 and base-Rpn14-Hsm3-Nas6(2) states, initial models were generated using two parts: 1. Rpt5, Rpt4, Rpt3, Rpt6, Nas6, Rpn14, Rpn2, and Rpn13; and 2. Rpt1, Rpt2, Hsm3, and Rpn1. For base-Hsm3-Nas6, initial models were generated using two parts: 1. Rpt5, Rpt4, Rpt3, Rpt6, Nas6, Rpn2, and Rpn13; and 2. Rpt1, Rpt2, Hsm3, and Rpn1. Initial models for local refinement maps were the same as for their parent maps. We observed that AlphaFold models were strongly biased toward producing structure models similar to a mature, fully closed base ring, which required significant further manual adjustment to build correct models into our densities. Structured parts of AlphaFold models were then separately fitted into the maps using ChimeraX ^51^, while allowing unstructured region to be flexible. The models were then manually inspected and adjusted using ISOLDE in ChimeraX based on observed densities ^52^. The models were finally refined using real-space refinement jobs in PHENIX ^53^ without simulated annealing and morphing, and validation statistics were also generated by PHENIX.

### Single-molecule fluorescence microscopy

PEGylated coverslips modified with biotin were prepared using established protocols ^54^. Just before the experiment, the flow chamber was built on the coverslip and neutravidin was immobilized on the coverslip through biotin. The Nas2 module containing biotinylated Rpt4 and Nas2-LD555 was pre-assembled with the Rpn14/Nas6 module at 1 µM each on ice for one hour. The preassembled modules were diluted to 100 pM and immobilized on to the coverslip. The Hsm3 module with Hsm3-LD655 was diluted to 10 nM, flowed onto the coverslip, and immediately monitored using TIRF microscopy. Briefly, we used a Nikon Ti2 TIRF microscope setup with Nikon LUN-F laser box, 60× oil objective (1.49 NA, Nikon), perfect focus system (Nikon), and filter cube TRF59907 (Chroma) ^38^. Movies were taken at 20 frames per s for 60 s on an iXon Ultra 897 (Andor) EMCCD camera with 532 nm and 640 nm laser simultaneous TIR excitation. The fluorescent signal was split into green and red channels using a Hamamatsu W-view Gemini equipped with a T640lpxr-UF2 dichroic mirror (Chroma) and ET595/50-m and ET655lp bandpass filters (Chroma). The movies were analyzed using iSMS software suite ^55^. Briefly, preassembled Nas2 and Rpn14/Nas6 modules were localized by picking green fluorophore spots in the green channel, and both green and red fluorescence time traces were extracted at the same position in their respective channels using a cross-channel alignment map obtained previously with TetraSpeck beads (Thermo Fisher Scientific). The time traces were manually inspected to find colocalization of the two fluorophores. Only eight time traces were determined confidently to show colocalization from six movies due to low association rate limited by the low concentration required for this single molecule method. The dwell times of different chaperones were extracted from these time traces using the transitions defined in Fig. 2B. To determine an estimate of the lifetime of these dwell events, we used a maximum likelihood estimation (MLE) method as described below. MLE of the expected value of lifetime assuming a single exponential distribution is given by: 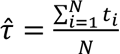, where *t_i_* is the i-th observed dwell time and N is the number of observed dwell time. MLE of the standard error of the lifetime is given by: 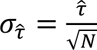. The values of lifetime reported in the main text were written as 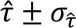.

## Resource availability

### Lead contact

Further information and requests for resources and reagents should be directed to and will be fulfilled by the lead contact, Andreas Martin.

### Materials availability

All constructs generated in this study are available from the lead contact upon request and completion of a Material Transfer Agreement.

### Data and Code Availability

- All data generated or analyzed during this study are included in this manuscript and the Supplementary materials. The cryo-EM density maps and corresponding atomic coordinates for the assembly intermediates of the yeast 26S proteasome base can be found on the Electron Microscopy Data Bank (EMDB) and Protein Data Bank (PDB) under the following accession codes: EMD-77338/PDB-36AX (base-Nas2-Rpn14-Hsm3-Nas6), EMD-77350/PDB-36BD (base-Rpn14-Hsm3-Nas6), EMD-77358/PDB-36BL (base-Rpn14-Hsm3-Nas6(2)), EMD-77359/PDB-36BM (base-Hsm3-Nas6), EMD-77307/PDB-35ZV (Rpn14-Rpt6 locally refined from base-Rpn14-Hsm3-Nas6(2)), EMD-77304 (Hsm3-Rpt1-Rpt2 locally refined from base-Rpn14-Hsm3-Nas6), EMD-77302/PDB-35ZR (Hsm3-Rpt1-Rpt2 locally refined from base-Hsm3-Nas6), EMD-77308/PDB-35ZW (Hsm3-Rpt1-Rpt2-Rpt3-Rpt4-Rpt5 locally refined from base-Hsm3-Nas6).
- This paper does not report original code.
- Any additional information required to reanalyze the data reported in this paper is available from the lead contact upon request.

## Acknowledgments

We thank all members of the Martin lab for discussion and support. Cryo-EM data were collected at the UCB Cal-Cryo facility, and we thank Kedar Sharma for cryo-EM operational support. Mass photometry data were acquired at UCB QB3 Cell and Tissue Analysis Facility, and we thank Mary West for support with the mass photometry instrument.

## Author contributions

H.H.H. and A.M. conceived the study and designed experiments, H.H.H. cloned constructs, expressed, and purified proteins, performed biochemical, single-molecule, and mass-photometry measurements, and did the data analyses. H.H.H. also performed cryo-EM sample preparation, data collection, and data processing, and generated atomic models based on cryo-EM maps. H.H.H. and A.M. wrote the manuscript.

## Extended Data

**Extended Data Figure 1.**
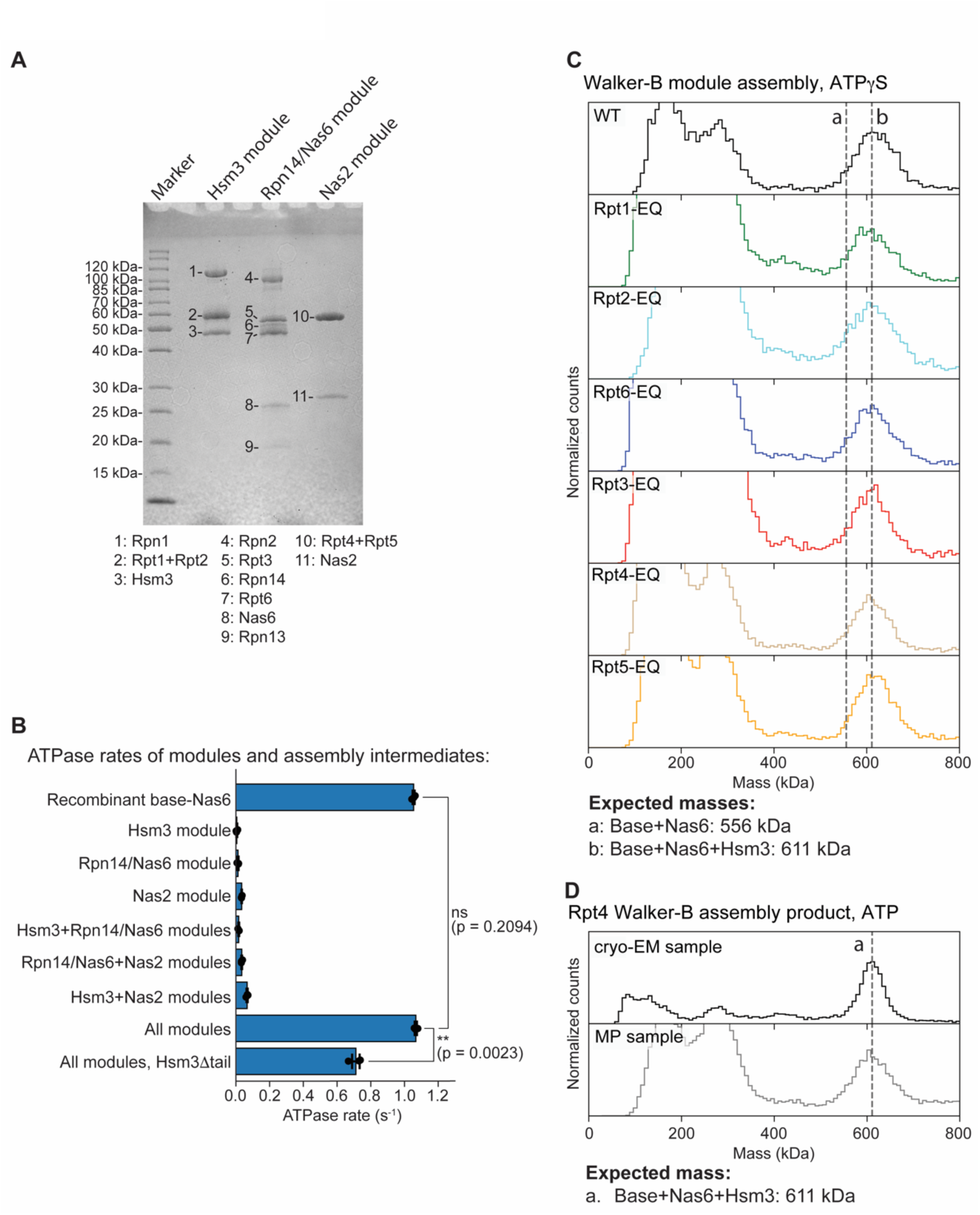
Reconstitution of the Hsm3, Rpn14/Nas6, and Nas2 modules, ATPase measurements, and mass-photometry analyses. (A) Coomassie-stained SDS-PAGE gel showing the components of the recombinantly expressed base-assembly modules. (B) ATP-hydrolysis measurement of individual modules, their combinations, and the recombinantly expressed Nas6-bound base as control. Shown are the mean values and standard deviations of the mean for three technical replicates. Statistical significance was calculated using a one-way ANOVA test: ns, p = 0.2094; **, p = 0.0023. (C) Mass-photometry analysis of the assembly products for modules with individual Walker-B EQ-mutant Rpt subunits in the presence of ATPγS. Assembly condition were similar to those shown in Fig. 1E, except for using ATPγS instead of ATP, which reveals that the difference observed for Rpt4-EQ compared to other Rpt-EQ variants in ATP were due to the lack of ATP hydrolysis in this subunit. (D) Mass-photometric comparison between the sample used for cryo-EM and a separately assembled base with Rpt4-EQ shown in Fig. 1E. No difference is observed for the mass peaks of the two measurements, indicating that the difference in Rpn14 and Nas2 occupancy between the cryo-EM structures and the mass-photometry measurement was due to the ∼ 2000-fold higher concentration.

**Extended Data Figure 2.**
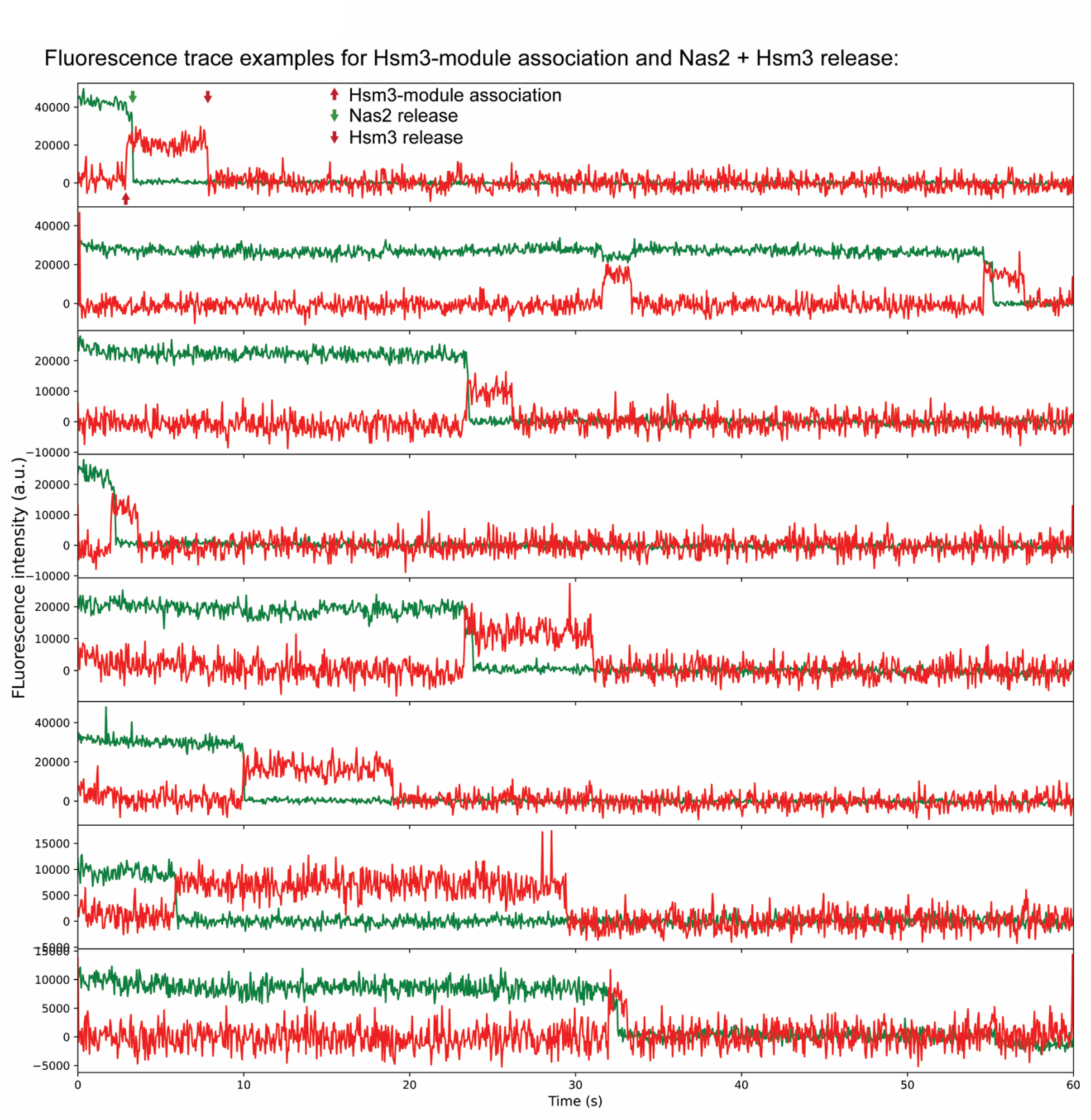
Example single-molecule fluorescence traces for module assembly and chaperone release. Conditions were identical to the measurement show in Fig. 2C (included here again in panel 5), with the LD655-labeled Hsm3 module associating with the immobilized Rpn14/Nas6 and LD555-Nas2 modules, and following the subsequent release of Hsm3 and Nas2 (indicated by red and green arrows).

**Extended Data Figure 3.**
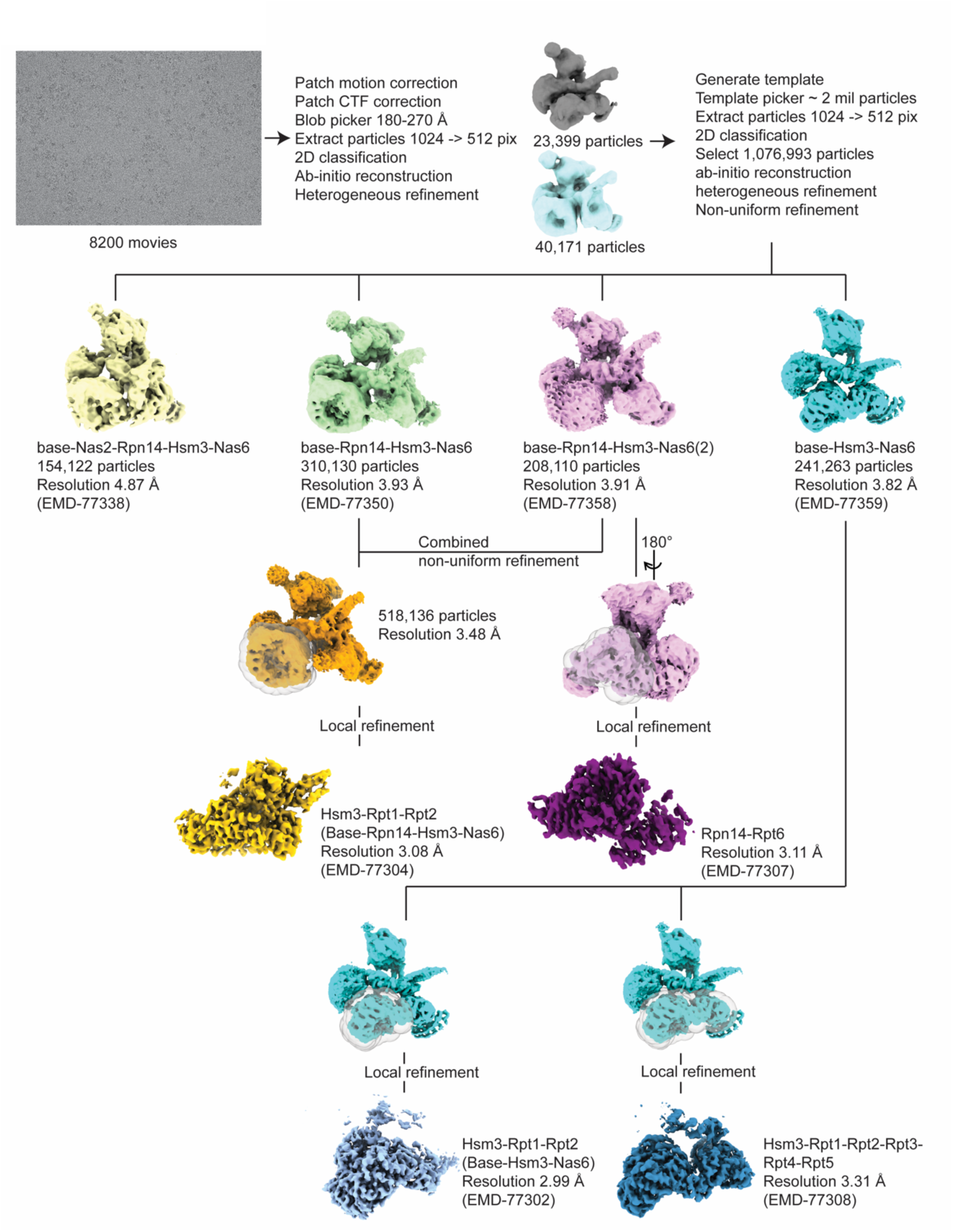
Workflow chart for the cryo-EM data processing.

**Extended Data Figure 4.**
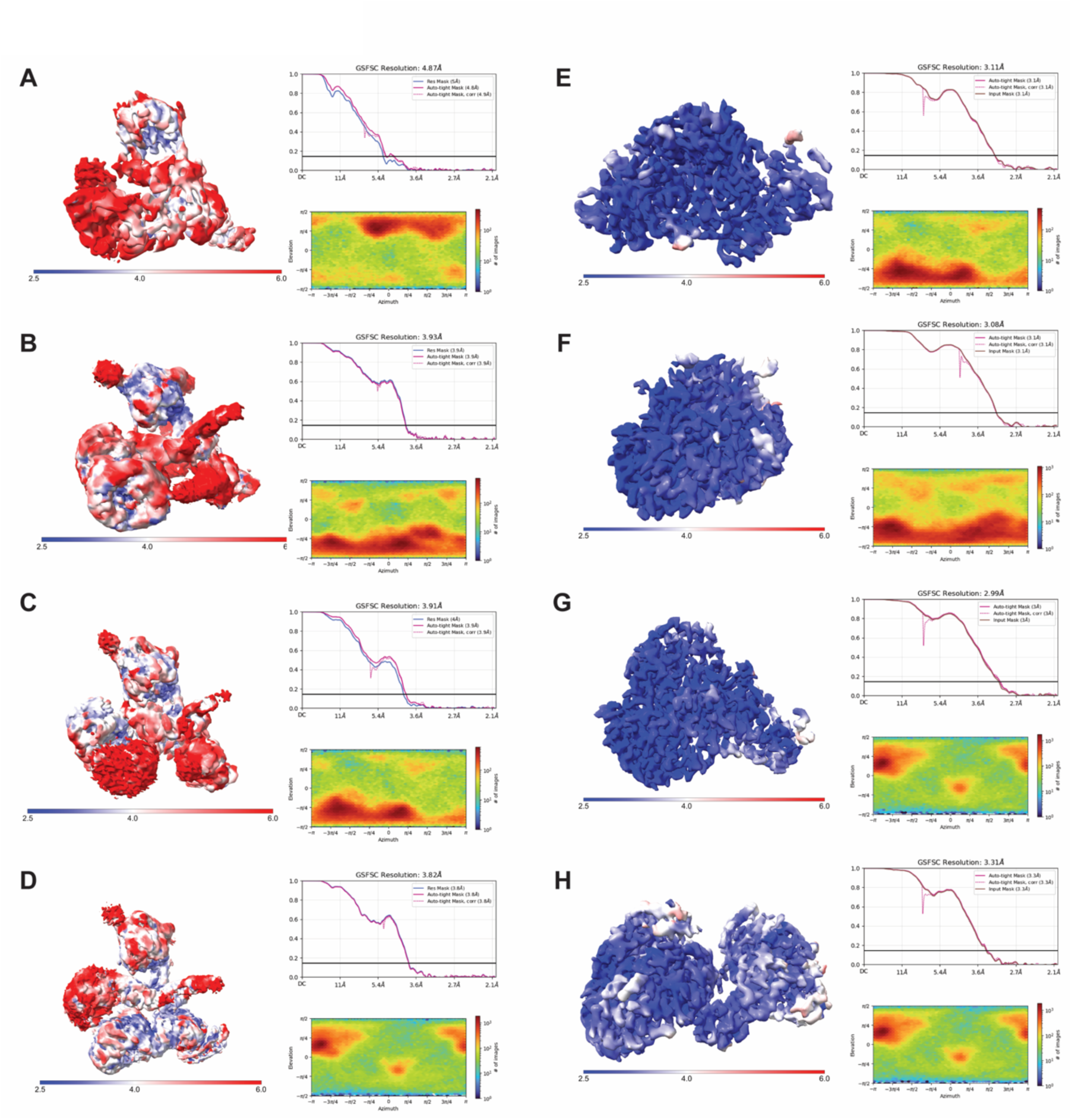
Local resolution and particle orientation distribution of cryo-EM density maps. (A-H) Color maps showing local resolution, gold-standard Fourier shell correlation plots, and particle orientation distributions for (A) base-Nas2-Rpn14-Hsm3-Nas6, (B) base-Rpn14-Hsm3-Nas6, (C) base-Rpn14-Hsm3-Nas6(2), (D) base-Hsm3-Nas6, (E) the Rpn14-Rpt6 local refinement, (F) the Hsm3-Rpt1-Rpt2 local refinement within base-Rpn14-Hsm3-Nas6, (G) the Hsm3-Rpt1-Rpt2 local refinement within base-Hsm3-Nas6, and (H) the Hsm3-Rpt1-Rpt2-Rpt3-Rpt4-Rpt5 local refinement within base-Hsm3-Nas6.

**Extended Data Figure 5.**
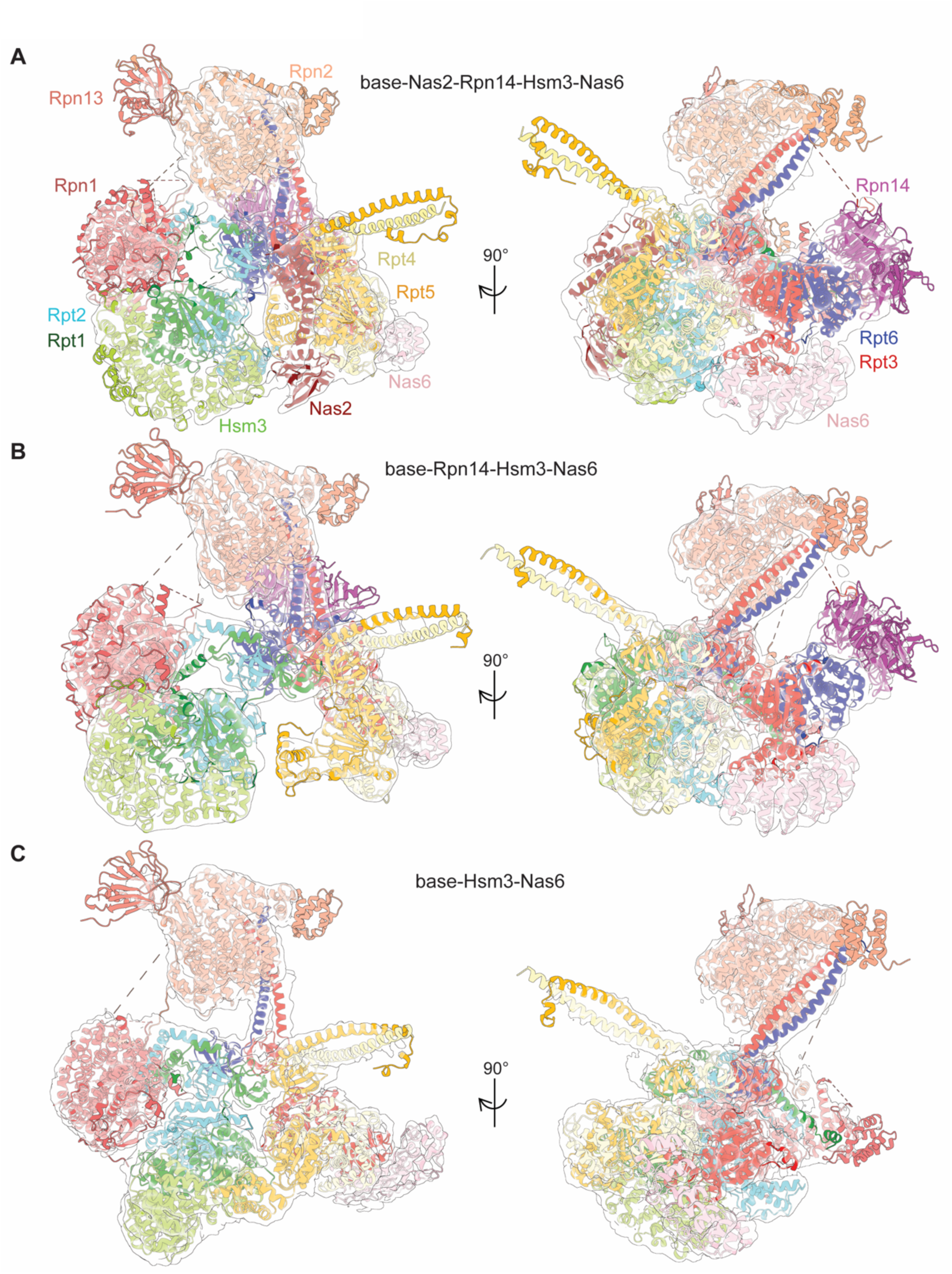
Molecular models and Cryo-EM maps of base assembly intermediates. (A-C) Molecular models (as shown in Fig. 3) are overlaid with cryo-EM densities for (A) base-Nas2-Rpn14-Hsm3-Nas6, (B) base-Rpn14-Hsm3-Nas6, and (C) base-Hsm3-Nas6 states.

**Extended Data Figure 6.**
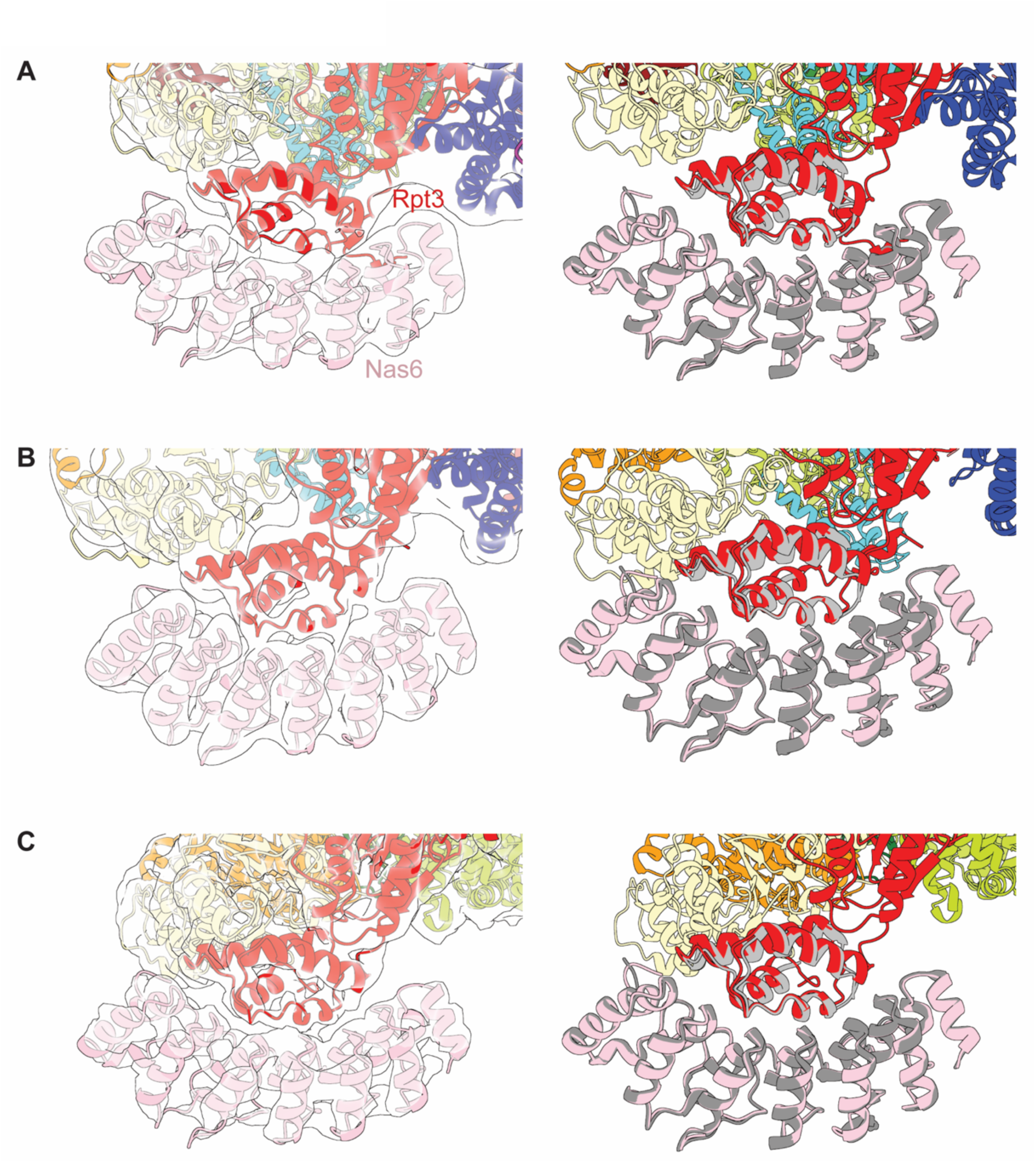
Nas6 interacts with the Rpt3 small AAA+ subdomain similarly across different assembly states. (A-C) Focused view of the Nas6-Rpt3 interaction for (A) base-Nas2-Rpn14-Hsm3-Nas6, (B) base-Rpn14-Hsm3-Nas6, and (C) base-Hsm3-Nas6 states. Left: Molecular models overlaid with cryo-EM densities, as in Extended Data Figure 5, but zoomed in on Nas6. Right: Crystal structure of the complex between Nas6 and the Rpt3 small AAA+ subdomain (PDB: 2DZN, grey) ^24^, aligned to Nas6 models from this study.

**Extended Data Figure 7.**
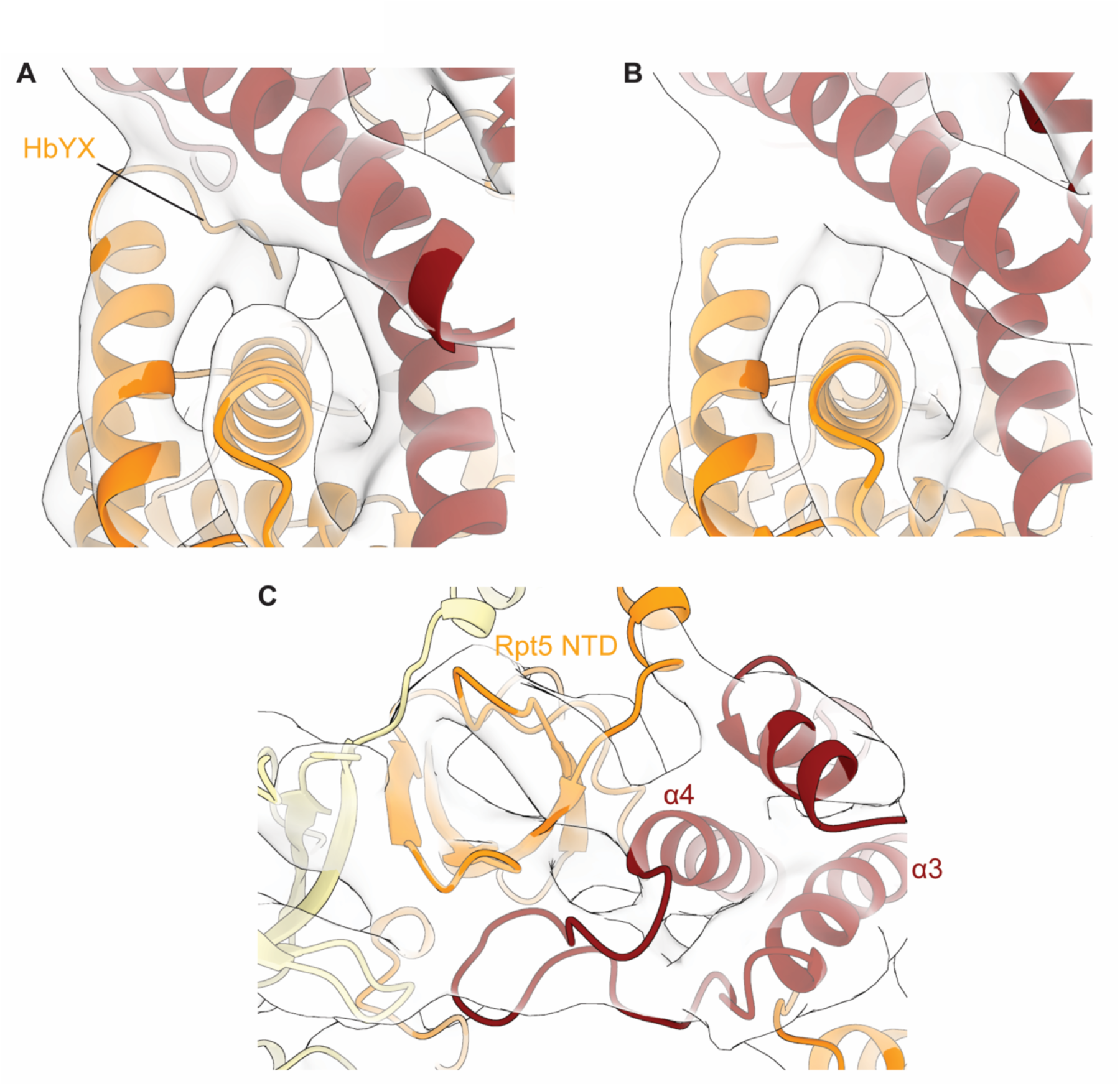
Nas2-Rpt5 interactions. Comparison of the Nas2-Rpt5 model from this study (A) with the previous crystal structure model (PDB:3WHL) ^17^ (B), both docked into the density for base-Nas2-Rpn14-Hsm3-Nas6. Additional density indicates that the Rpt5’s C-terminal HbYX motif is sandwiched between Nas2’s N-terminal helices and Rpt5’s small AAA+ subdomain (A). (C) Closed-up view of Nas2 interaction with Rpt5’s NTD. Note that the loop connecting Nas2’s N-terminal helices α3 and α4 also interacts with Rpt5’s NTD.

**Extended Data Figure 8.**
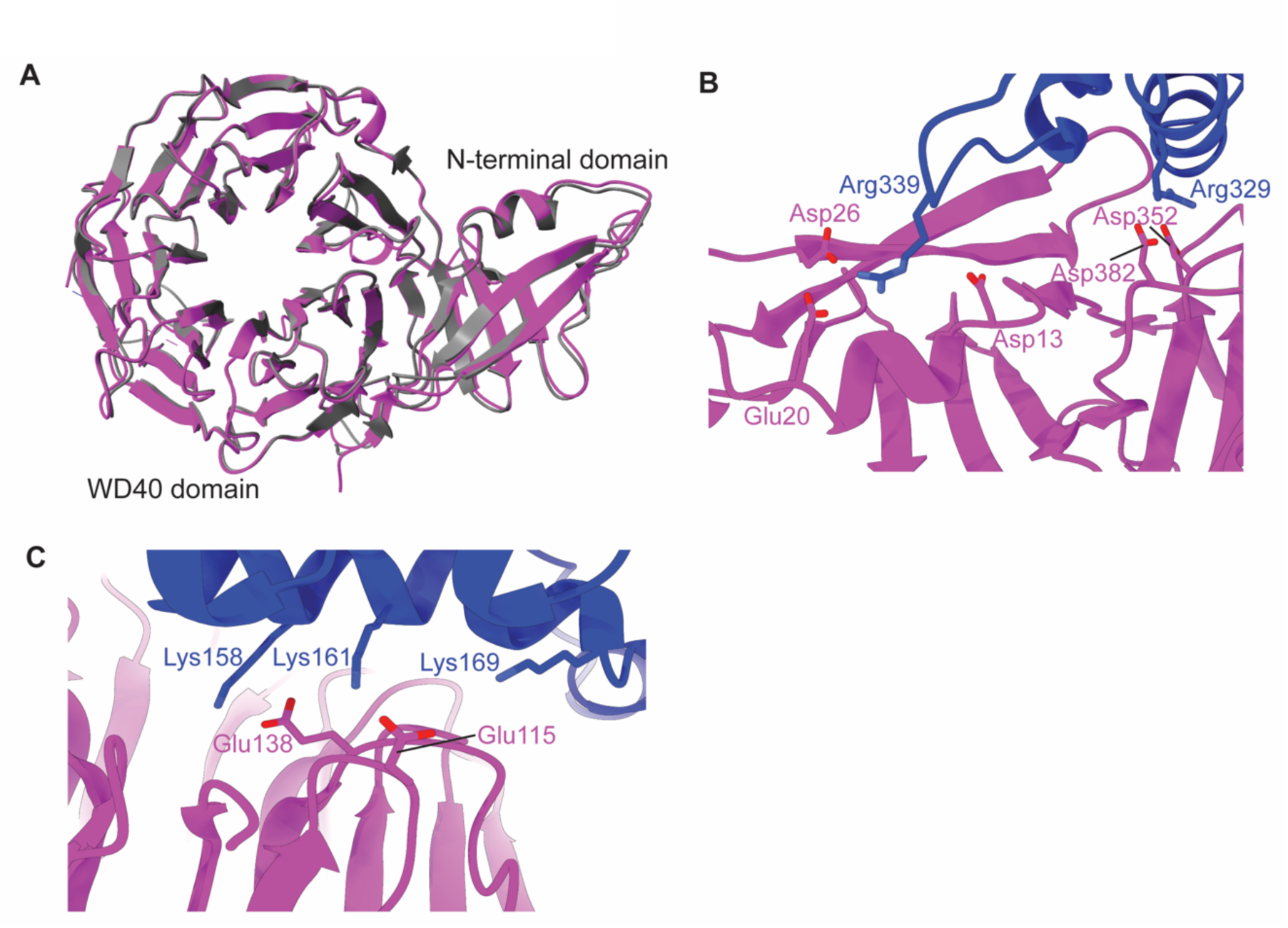
Rpn14-Rpt6 interactions. (A) Our structure of the Rpt6-bound Rpn14 (magenta) is similar to the previous crystal structure of isolated Rpn14 (PDB:3ACP, grey) ^30^. (B) Multiple negatively charged residues in Rpn14 N-terminal domain (Asp13, Glu20, Asp26, Asp352 and Asp382) form interface with multiple positively charged residues of Rpt6 small AAA+ subdomain (Arg329 and Arg339). (C) Positively charged residues of Rpt6 ATPase large domain (Lys158, Lys161 and Lys169) interact with a patch of negatively charged residues of Rpn14 WD40 domain (Glu115 and Glu138)

**Extended Data Figure 9.**
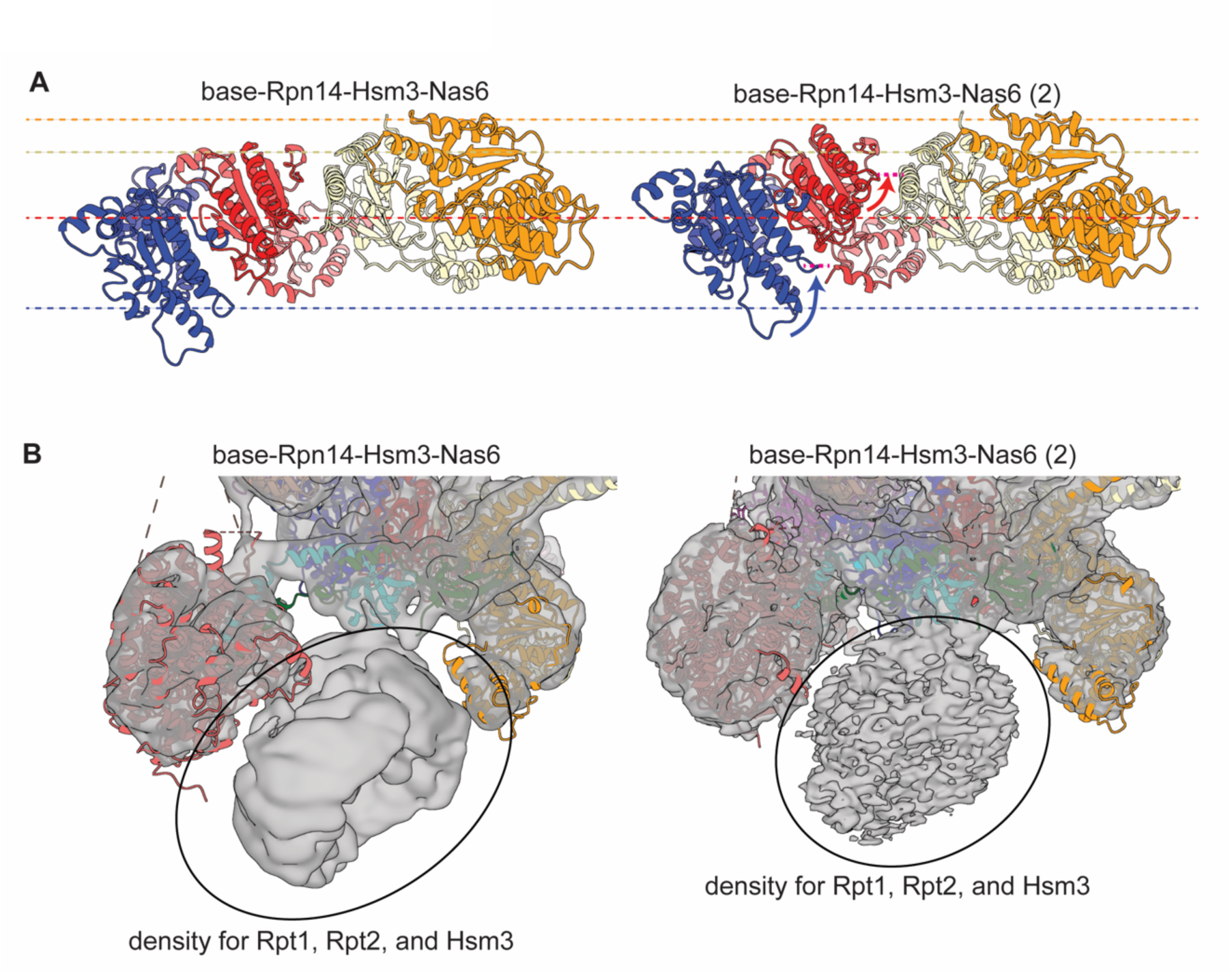
Rpn14-Rpn1 interaction allows rearrangement of Rpt ATPases. (A) Comparison of the ATPase-domain positions in the AAA+ ring plane shows that Rpt6 and Rpt3 tilt up and become more planar with Rpt4 and Rpt5 as Rpn14 and Rpn1 interact during the transition from the base-Rpn14-Hsm3-Nas6 (left) to the base-Rpn14-Hsm3-Nas6(2) conformation (right). Horizontal lines in corresponding colors are drawn from the pore-1 loop of each Rpt in the base-Rpn14-Hsm3-Nas6 conformation to show the relative movement of Rpts during the transition, with the new positions of Rpt6’s and Rpt3’s pore-1 loops indicate by magenta lines. (B) Density of the Hsm3-Rpt1-Rpt2 portion shows increased flexibility as Rpn14 and Rpn1 make contact. Molecular models of Hsm3 and the ATPase domains of Rpt1 and Rpt2 are not shown to better visualize their corresponding densities.

**Extended Data Figure 10.**
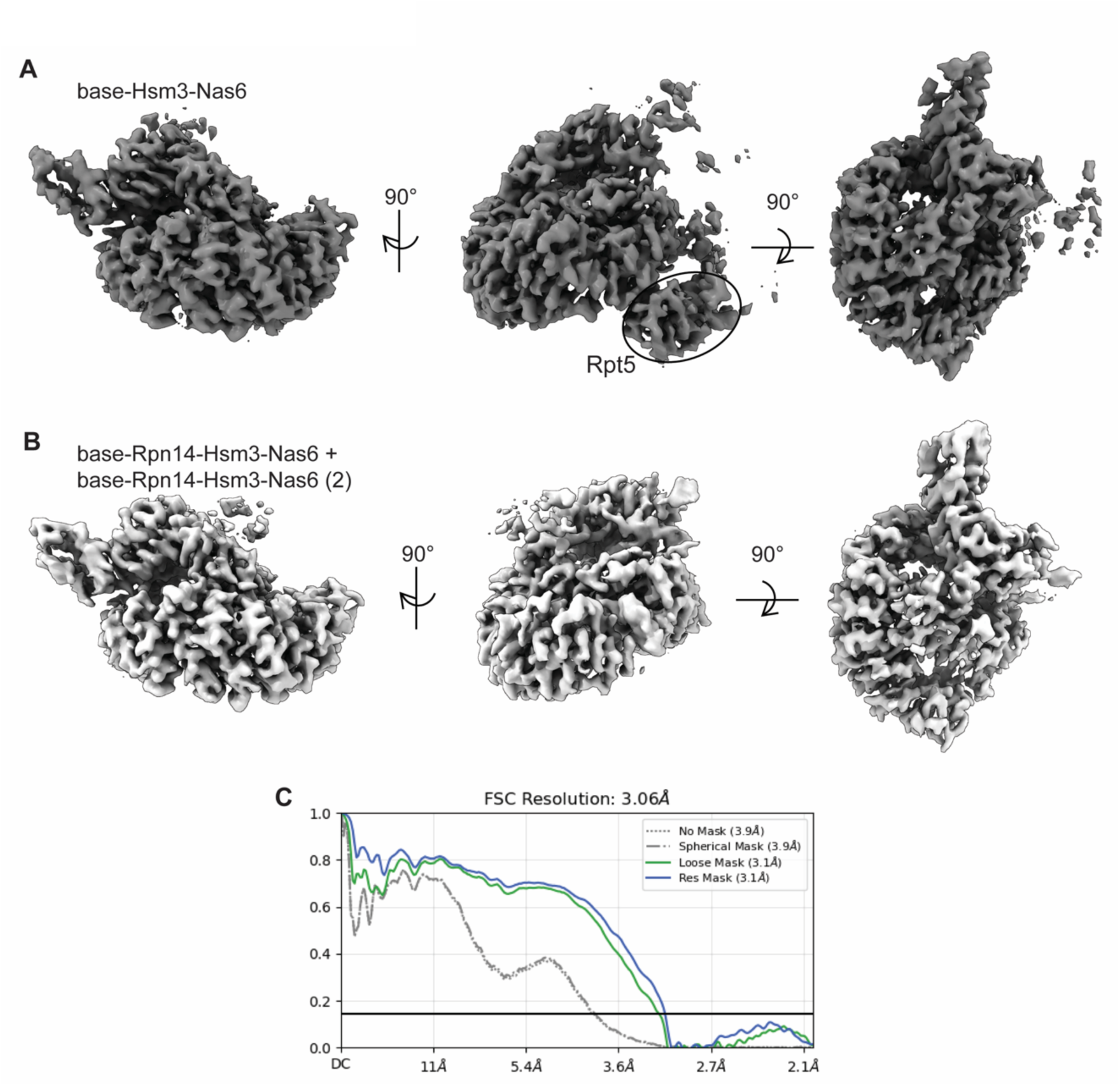
Hsm3 and the ATPase domains of Rpt1 and Rpt2 adopt the same structure in the base-Rpn14-Hsm3-Nas6 and base-Hsm3-Nas6 states. (A, B) Local-refinement density maps for the Hsm3-bound ATPase domains of Rpt1 and Rpt2 in (A) base-Hsm3-Nas6 and (B) base-Rpn14-Hsm3-Nas6 + base-Rpn14-Hsm3-Nas6(2) particles. (C) Fourier shell correlation between the density maps from (A) and (B).

**Extended Data Figure 11.**
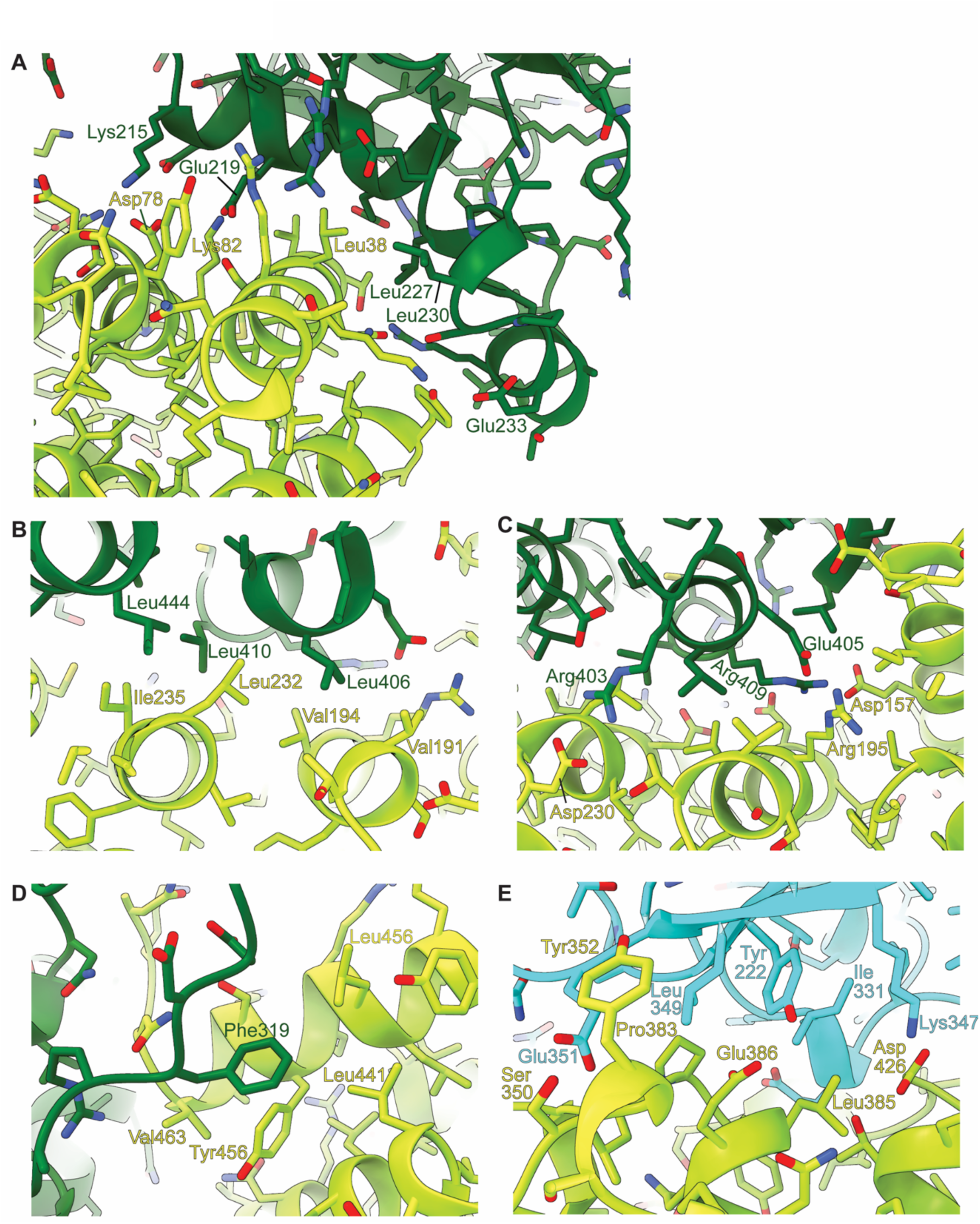
Hsm3 interactions with the ATPase domains of Rpt1 and Rpt2. (A) Rpt1’s helix 214-239 forms mixed hydrophilic and hydrophobic interactions with Hsm3’s N-terminal helices 1-86. (B) Hydrophobic core formed between Hsm3 and Rpt1’s small AAA+ subdomain (Hsm3: Val191, Val194, Leu232 and Ile235; Rpt1: Leu406, Leu410 and Leu444). (C) Charge-charge interactions between Hsm3 and Rpt1’s small AAA+ subdomain (Hsm3-Rpt1: Glu157-Arg409, Arg195-Glu405 and Asp230-Arg403) surrounding the hydrophobic core shown in (B). (D) Hsm3 forms hydrophobic interaction (Leu441, Leu456, Tyr459 and Val463) with Phe319 in the pore-2 loop of Rpt1. (E) Hsm3-Rpt2 interaction involving hydrophobic interactions (Hsm2: Tyr352, Pro383 and Leu385; Rpt2: Tyr222, Ile331 and Leu349), potential hydrogen bonding (Hsm3-Rpt2: Ser350-Glu351 and Glu386-Tyr222), and charge-charge interaction (Hsm3-Rpt2: Asp426-Lys347).

**Extended Data Figure 12.**
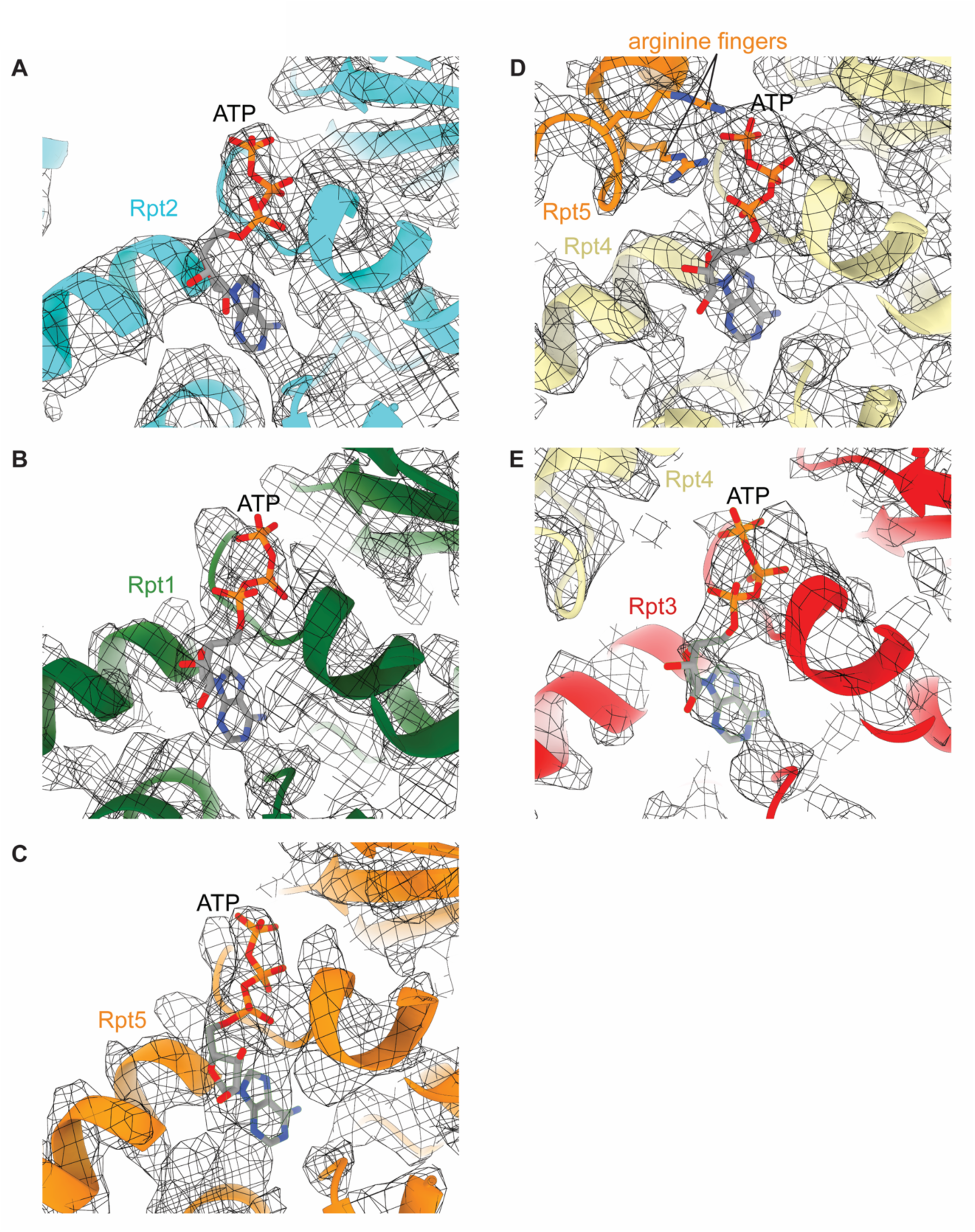
Nucleotide occupancies in the base-Hsm3-Nas6 state. (A-E) Structural models showing ATP bound to the ATPase site of (A) Rpt2, (B) Rpt1, (C) Rpt5, (D) Rpt4, and (E) Rp5. The ATPase sites are open in Rpt1, Rpt2, Rpt3, and Rpt5, and their ATP is not engaged by the neighboring subunit, whereas Rpt4’s site is closed and its ATP is contacted by the arginine fingers of Rpt5. Due to its high flexibility Rpt6 is not model in the base-Hsm3-Nas6 state and therefore not shown.

**Extended Data Figure 13.**
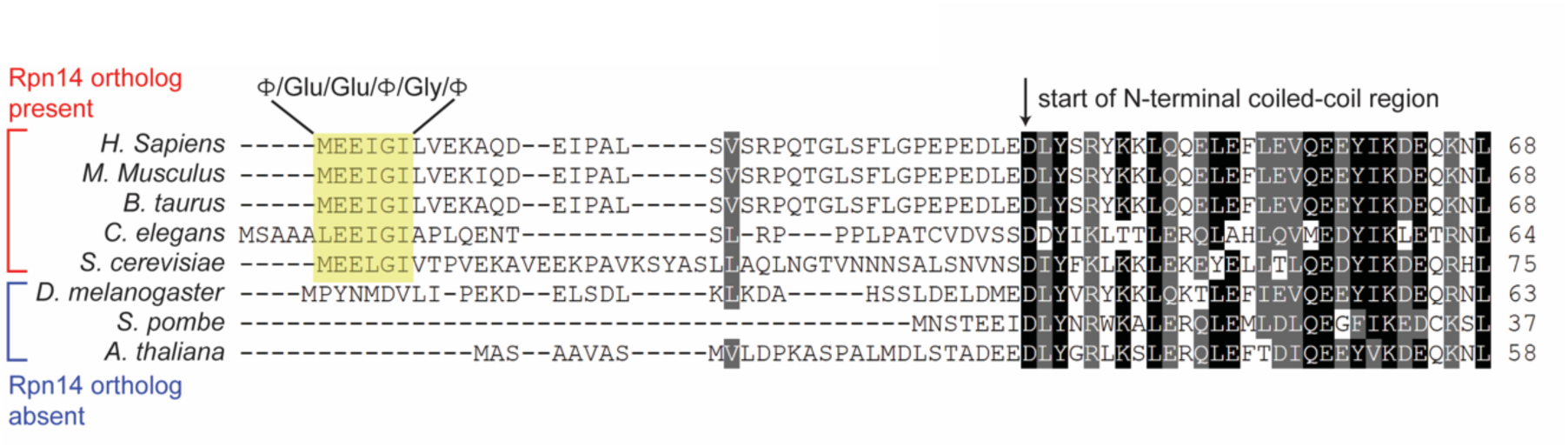
Multiple sequence alignment of Rpt3 from species with and without orthologs of Rpn14. Rpt3 sequences from species with Rpn14 orthologs show a conservation of the </Glu/Glu/</Gly/< motif (<: hydrophobic residue) that interacts with Rpn14’s WD40 domain. Species that lack a Rpn14 ortholog show no conservation of this motif in Rpt3.

---

**Supplemental Video 1:** Molecular model of the base-Nas2-Rpn14-Hsm3-Nas6 state, with Rpt3 (red), Rpt6 (blue), Rpt2 (cyan), Rpt1 (dark green), Rpt5 (orange), Rpt4 (tan), Rpn1 (Indian), Rpn2 (dark salmon), and Rpn13 (salmon) shown in surface representation, and the chaperones Nas6 (pink), Hsm3 (lime green), Nas2 (dark red), and Rpn14 (magenta) in cartoon representation.

**Supplemental Video 2:** Molecular model of the base-Nas2-Rpn14-Hsm3-Nas6 state in cartoon representation, focusing on the interaction between Nas6 (pink) and the small AAA+ subdomain of Rpt3 (red).

**Supplemental Video 3:** Molecular model of the base-Nas2-Rpn14-Hsm3-Nas6 state in cartoon representation, focusing on the interaction between Nas2 (dark red) and Rpt5 (orange), and on Nas2’s N-terminal domain as part of the N-ring of Rpt subunits. The NTD of Rpt1 (dark green) was not resolved in this state and is modeled in an approximate position outside the N-ring.

**Supplemental Video 4:** Molecular model of the base-Nas2-Rpn14-Hsm3-Nas6 state in cartoon representation, focusing on the interaction between Rpn14 (magenta) and Rpt6 (blue), with Rpn14’s N-terminal domain contacting the small AAA+ subdomain of Rpt6, while Rpn14’s WD propeller primarily interacts with the large AAA+ subdomain.

**Supplemental Video 5:** Morph showing the transition between base-Rpn14-Hsm3-Nas6 and base-Rpn14-Hsm3-Nas6(2), during which the Rpt6-bound Rpn14 forms contacts with Rpn1 to close a ∼ 60 Å gap.

**Supplemental Video 6:** Molecular model of the base-Nas2-Rpn14-Hsm3-Nas6 state in cartoon representation, focusing on the interaction between Hsm3 (lime green), Rpt1 (dark green), and Rpt2 (cyan).

**Supplemental Video 7:** Morph for the transition between base-Nas2-Rpn14-Hsm3-Nas6 and base-Rpn14-Hsm3-Nas6, showing the conformational changes that are coupled to the dissociation of Nas2 (dark red). In base-Nas2-Rpn14-Hsm3-Nas6, the NTD of Rpt1 (dark green) is not resolved and therefore modeled near the N-ring. Upon Nas2 dissociation, this NTD gets incorporated and completes the N-ring, while the ATPase domains of the six Rpts become even more splayed out.

**Supplemental Video 8:** Side view of the morph between base-Rpn14-Hsm3-Nas6(2) and base-Hsm3-Nas6, showing the conformational transitions, in particular the partial ATPase ring closure, that are coupled to the dissociation of Rpn14 (magenta). The ATPase domain of Rpt6 (blue) becomes more flexible and poorly resolved, and is therefore shown in a modeled position for the final base-Hsm3-Nas6 state.

**Supplemental Video 9:** Bottom view of the morph between base-Rpn14-Hsm3-Nas6(2) and base-Hsm3-Nas6, showing the conformational transitions that are coupled with the dissociation of Rpn14 (magenta). especially the rotation of the Rpt1/Rpt2 ATPase domains and the resulting partial closure of the AAA+ ring. The ATPase domain of Rpt6 (blue) becomes more flexible and poorly resolved, and is therefore shown in a modeled position for the final base-Hsm3-Nas6 state.

**Supplemental Video 10:** Model of the base-Hsm3-Nas6 state viewed from the bottom, with the Rpt and Rpn subunits initially shown in surface representation, while Hsm3 and Nas6 are shown as cartoons. The video then zooms in onto the central channel of the ATPase ring that is occupied by the C-terminal tail of Hsm3, mimicking an engaged substrate polypeptide. Further zooming in, rotating to a side view of the central channel, and removing Rpt1, Rpt2, and Rpt6 reveals how Rpt3 (red), Rpt4 (tan), and Rpt5 (orange) interact through their Tyr-containing pore-1 loops (magenta) in a spiral staircase conformation with Hsm3’s tail (lime green).

**Supplemental Video 11:** Morph for the transitions from base-Nas2-Rpn14-Hsm3-Nas6 via base-Rpn14-Hsm3-Nas6 and base-Rpn14-Hsm3-Nas6(2) to base-Hsm3-Nas6 as viewed from the bottom. It shows the conformational changes of the ATPase domains and the gradual AAA+ ring closure during the ejection of Nas2 and Rpn14.

**Supplemental Table 1.**

|  | base-Nas2-Rpn14-Hsm3-Nas6 | base-Rpn14-Hsm3-Nas6 | base-Rpn14-Hsm3-Nas6(2) | base-Hsm3-Nas6 | Rpn14-Rpt6<br>(base-Rpn14-Hsm3-Nas6(2)) | Hsm3-Rpt1-Rpt2 (base-Rpn14-Hsm3-Nas6) | Hsm3-Rpt1-Rpt2 (base-Hsm3-Nas6) | Hsm3-Rpt1-Rpt2-Rpt3-Rpt4-Rpt5 |
| --- | --- | --- | --- | --- | --- | --- | --- | --- |
| EM Data Bank | EMD-77338 | EMD-77350 | EMD-77358 | EMD-77359 | EMD-77307 | EMD-77304 | EMD-77302 | EMD-77308 |
| PDB | 36AX | 36BD | 36BL | 36BM | 35ZV | N/A | 35ZR | 35ZW |
| **Data collection and**<br>**processing** |  |  |  |  |  |  |  |  |
| Magnification | ×81000 | ×81000 | ×81000 | ×81000 | ×81000 | ×81000 | ×81000 | ×81000 |
| Voltage (kV) | 300 | 300 | 300 | 300 | 300 | 300 | 300 | 300 |
| Electron exposure (e-/Å<sup>2</sup>) | 50 | 50 | 50 | 50 | 50 | 50 | 50 | 50 |
| Defocus range (µm) | -0.5 to -2.0 | -0.5 to -2.0 | -0.5 to -2.0 | -0.5 to -2.0 | -0.5 to -2.0 | -0.5 to -2.0 | -0.5 to -2.0 | -0.5 to -2.0 |
| Pixel size (Å) | 1.048 | 1.048 | 1.048 | 1.048 | 1.048 | 1.048 | 1.048 | 1.048 |
| Symmetry imposed | C<sub>1</sub> | C<sub>1</sub> | C<sub>1</sub> | C<sub>1</sub> | C<sub>1</sub> | C<sub>1</sub> | C<sub>1</sub> | C<sub>1</sub> |
| Initial particle images (no.) | 1076993 | 1076993 | 1076993 | 1076993 | 1076993 | 1076993 | 1076993 | 1076993 |
| Final particle images (no.) | 154122 | 310130 | 208110 | 241263 | 208110 | 518136 | 241263 | 241263 |
| Map resolution (Å) | 4.87<br>0.143 | 3.93<br>0.143 | 3.91<br>0.143 | 3.82<br>0.143 | 3.11<br>0.143 | 3.08<br>0.143 | 2.99<br>0.143 | 3.31<br>0.143 |
| FSC threshold | 0.143 | 0.143 | 0.143 | 0.143 | 0.143 | 0.143 | 0.143 | 0.143 |
| **Refinement** |  |  |  |  |  | no model<br>built |  |  |
| Model resolution (Å) | 5.02 | 3.89 | 3.96 | 3.79 | 3.04 |  | 2.95 | 3.26 |
| FSC threshold | 0.143 | 0.143 | 0.143 | 0.143 | 0.143 |  | 0.143 | 0.143 |
| Map sharpening *B* factor<br>(Å<sup>2</sup>) | NA | NA | NA | NA | NA |  | NA | NA |
| Model composition |  |  |  |  |  |  |  |  |
| Non-hydrogen atoms | 43182 | 41629 | 41530 | 36449 | 6397 |  | 7786 | 15764 |
| Protein residues | 5472 | 5280 | 5266 | 4617 | 807 |  | 971 | 1979 |
| Ligands | ATP: 6 | ATP: 6 | ATP: 6 | ATP: 5 | ATP: 1 |  | ATP: 2 | ATP: 5 |
| *B* factors (Å<sup>2</sup>)<br>(min/max/mean) |  |  |  |  |  |  |  |  |
| Protein | 149.33/1048.92/<br>615.27 | 79.32/779.92/31<br>7.85 | 88.90/1028.38/3<br>66.6 | 45.22/592.47/2<br>21.15 | 58.46/171.25/<br>102.06 |  | 63.19/208.03/<br>108.01 | 31.60/533.4<br>1/154.35 |
| Ligand | 247.85/569.79/4<br>04.63 | 201.25/515.45/3<br>35.82 | 189.94/712.02/4<br>19.10 | 124.80/187.07/<br>143.52 | 91.43/127.62/<br>102.87 |  | 82.62/135.43/<br>100.03 | 47.76/170.8<br>4/105.23 |
| R.m.s. deviations |  |  |  |  |  |  |  |  |
| Bond lengths (Å) | 0.004 | 0.004 | 0.003 | 0.003 | 0.003 |  | 0.004 | 0.003 |
| Bond angles (°) | 0.706 | 0.664 | 0.659 | 0.676 | 0.64 |  | 0.664 | 0.69 |
| Validation |  |  |  |  |  |  |  |  |
| MolProbity score | 2 | 1.71 | 1.57 | 1.75 | 1.49 |  | 1.68 | 1.59 |
| Clashscore | 17.55 | 11.21 | 9.29 | 12 | 6.83 |  | 8.42 | 9.66 |
| Poor rotamers (%) | 0 | 0.85 | 0.92 | 1.27 | 1.27 |  | 1.97 | 1.05 |
| Ramachandran plot |  |  |  |  |  |  |  |  |
| Disallowed (%) | 0 | 0.02 | 0.06 | 0 | 0 |  | 0 | 0 |
| Allowed (%) | 3.83 | 2.75 | 2.3 | 2.36 | 2.16 |  | 1.77 | 2.3 |
| Favored (%) | 96.17 | 97.23 | 97.65 | 97.64 | 97.84 |  | 98.23 | 97.7 |

